# Memory-specific encoding activities of the ventral tegmental area dopamine and GABA neurons

**DOI:** 10.1101/2023.06.28.546967

**Authors:** Vasileios Glykos, Shigeyoshi Fujisawa

## Abstract

Although the midbrain dopamine (DA) system plays a crucial role in higher cognitive functions, including updating and maintaining short-term memory, the encoding properties of the somatic spiking activity of ventral tegmental area (VTA) DA neurons for short-term memory computations have not yet been identified. Here, we probed and analyzed the activity of optogenetically identified DA and GABA neurons while mice engaged in short-term memory-dependent behavior in a T-maze task. Single-neuron analysis revealed that significant subpopulations of DA and GABA neurons responded differently between left and right trials in the memory delay. With a series of control behavioral tasks and regression analysis tools, we show that firing rate differences are linked to short-term memory-dependent decisions and cannot be explained by reward-related processes, motivated behavior, or motor-related activities. This evidence provides novel insights into the mnemonic encoding activities of midbrain DA and GABA neurons.

## Introduction

Dopamine (DA) neurons originating in the ventral tegmental area (VTA) project to diverse forebrain regions, forming distinct but interacting neuromodulatory systems that are thought to play pivotal roles in the regulation of reward-related learning, motivation, and cognition (Sawaguchi and Goldman-Rakic, 1991; Schultz et al., 1993; Goldman-Rakic, 1995; Schultz et al., 1997; Tzschentke, 2001; Schultz, 2002; Pierce and Kumaresan, 2006; Berridge, 2007; Vijayraghavan et al., 2007; Lammel et al., 2008; Robbins and Arnsten, 2009; Hauber, 2010; Cohen et al., 2012; Salamone and Correa, 2012; Howe et al., 2013; Matsumoto and Takada, 2013; Hamid et al., 2016; Mohebi et al., 2019). A wealth of electrophysiological recordings from midbrain DA neurons, complemented by *in vivo* microdialysis data indicate that midbrain DA activity promotes behaviors associated with motivation (Wise, 2004; Berridge, 2007; Salamone and Correa, 2012; Howe et al., 2013; Matsumoto and Takada, 2013; Hamid et al., 2016; Mohebi et al., 2019) and supports reward-based learning by encoding reward prediction error (RPE) signals (Schultz et al., 1993; Schultz et al., 1997; Cohen et al., 2012).

Also, DA is of central importance to higher cognitive functions, such as updating and maintaining short-term memory (Sawaguchi and Goldman-Rakic, 1991; Miller and Cohen, 2001; Ott and Nieder, 2019). Pioneering behavioral studies which pharmacologically manipulated the activity of DA receptors in the PFC revealed the significant role of DA signals on short-term memory. In fact, an inverted-U-shape effect was discovered, where too little or too much DA receptor stimulation impairs PFC-engaging short-term memory (Sawaguchi and Goldman-Rakic, 1991; Vijayraghavan et al., 2007; Robbins and Arnsten, 2009). Moreover, at the origin of the DA system, electrophysiological recordings at the VTA showed that DA neurons are not active in the delay period of memory tasks (Schultz et al., 1993; Schultz, 2002; Phillips et al., 2004; Matsumoto and Takada, 2013; Choi et al., 2020).

Motivated by the response of DA neurons to reward-related stimuli and memory delays, several lines of computational modeling studies sought to answer when and how DA signals support short-term memory “update” and “maintenance”. They proposed the “gating theory”, which provided a unified computational framework for reward prediction and short-term memory (Cohen et al., 2002; Dreher and Burnod, 2002; Montague et al., 2004; Ott and Nieder, 2019). According to the model, reward-predicting cues, elicit phasic DA release which opens the gate for the afferent signals to be stored in memory (update). But, in the delay period, low, tonic DA levels close the gate for interfering signals to enter the PFC and overwrite the short-term memory component (maintenance). Although the “gating theory” fits adequately the behavior-unique responses of DA neurons to the coding schemes of short-term memory, it relies mainly upon empirical evidence of putative DA neurons and the longstanding consensus that short-term memory depends on the unbroken chain of persistent neuronal activity (Durstewitz et al., 2000; Curtis and D’Esposito, 2003).

However, recent advances in the study of the brain’s functional organization suggest that persistent neuronal activity might not be the only candidate mechanism for the active maintenance of goal representation over short delays, leading to the proposal of new coding schemes for short-term memory (Stokes, 2015; Miller et al., 2018). One of these candidate mechanisms regards the memory-dependent dynamic changes in functional connectivity. Neural oscillations are abundant in the mammalian brain and are thought to offer the networking framework for the temporal organization of neuronal activity and information processing in short-term memory (Uhlhaas and Singer, 2006; Buschman et al., 2012; Miller et al., 2018). Calculating the phase coherence of neural oscillations between distributed brain regions, provides an estimation of the functional connectivity between them (Fries, 2005). Among other basal ganglia regions, the VTA engages dynamically in the large-scale network of brain systems that support memory-related information processing. Simultaneous electrophysiological recordings were performed in the PFC and the VTA while rodents executed memory-guided behavioral choices in a T-maze task (Fujisawa and Buzsáki, 2011). Neural oscillations (4Hz) were prominent in both regions throughout the task, but their power and coherence were adaptively increased in memory delay. In a similar behavioral task, another short-term memory-related coding scheme was reported, this time at the single neuronal level. It was shown that while rodents navigate the maze, performing memory-guided decisions, PFC and parietal neurons differentiate their firing activities between opposite behavioral choices (Fujisawa et al., 2008; Harvey et al., 2012). To summarize, this novel empirical evidence from rodent studies on the T-maze behavioral apparatus complements the coding framework of short-term memory with more dynamic and adaptive information-processing mechanisms other than persistent activity.

In studying the role of DA neurons in short-term memory, we should take into consideration that the DA neuronal circuit is by no means self-contained and therefore it should not be investigated in isolation. Neurons utilizing GABA as a neurotransmitter constitute approximately 30% of the VTA neuronal population. The memory-related encoding properties of these inhibitory neurons have been largely overlooked, despite evidence of a strong inhibitory influence on neighboring DA neurons (Nair-Roberts et al., 2008; Omelchenko and Sesack, 2009; Tan et al., 2012; van Zessen et al., 2012) and well-established interconnections with the PFC circuit (Carr and Sesack, 2000a, b).

In light of the above, we wished to investigate with fine temporal and spatial resolution the firing activity of optogenetically identified DA and GABA neurons while mice performed a T-maze reward-seeking task with memory load. We took into consideration that (i) earlier studies analyzed either the activity of putative DA neurons or drew inferences of the population activity from voltammetry and fiber photometry recordings (Schultz et al., 1993; Schultz, 2002; Phillips et al., 2004; Matsumoto and Takada, 2013; Choi et al., 2020), and (ii) field potentials (like the 4Hz oscillations recorded in the VTA) stem mainly from phase-aligned excitatory or inhibitory post-synaptic potentials, whereas spiking activity is sparse (Traub et al., 2004; Buzsaki, 2006). (iii) A recent report revealed the causal relationship of DA activity with short-term memory, by inhibiting DA neurons with optogenetic tools. However, they did not report the encoding properties of single VTA DA neurons (Choi et al., 2020).

## Results

### Optogenetic identification of DA and GABA neurons in the VTA

In the present study, we sought to investigate the encoding properties of DA and GABA neurons of the VTA while mice engage in memory-dependent reward-seeking behavior. To identify neurons, we expressed the light-gated cation channel, channelrhodopsin-2 (ChR2), in DA and GABA neurons by injecting an adeno-associated virus containing FLEX-ChR2 into DAT-Cre and VGAT-Cre transgenic mice, respectively (Bäckman et al., 2006; Tsai et al., 2009; Vong et al., 2011) (Figure 1A). Optogenetic identification and parallel electrophysiological recordings were performed using a custom-made diode-probe system (diode-fiber assemblies attached to high-density silicon probes (Stark et al., 2012), (Figures 1A and S1). For each neuron, we assessed the response to light pulse trains delivered before and after behavioral sessions (Figures 1, S1 and S2). We identified 104 neurons recorded from five DAT-Cre mice (hereafter referred to as DA neurons) and 74 neurons recorded from four VGAT-Cre mice (GABA neurons) with significant excitatory responses to light pulses (Figure 1B). Light-induced spikes from these neurons were almost identical to spontaneous spikes (waveform correlation coefficient > 0.9, Figure S1E and 1F). In addition, the electrophysiological profiles of the identified neuronal populations resembled those of previous studies (i.e., DA neurons fired action potentials with both wider waveforms and slower spontaneous firing rates than GABA neurons; Figure S1G), confirming the selective expression of ChR2 in DA and GABA neurons (Cohen et al., 2012; Tan et al., 2012).

**Figure 1.**
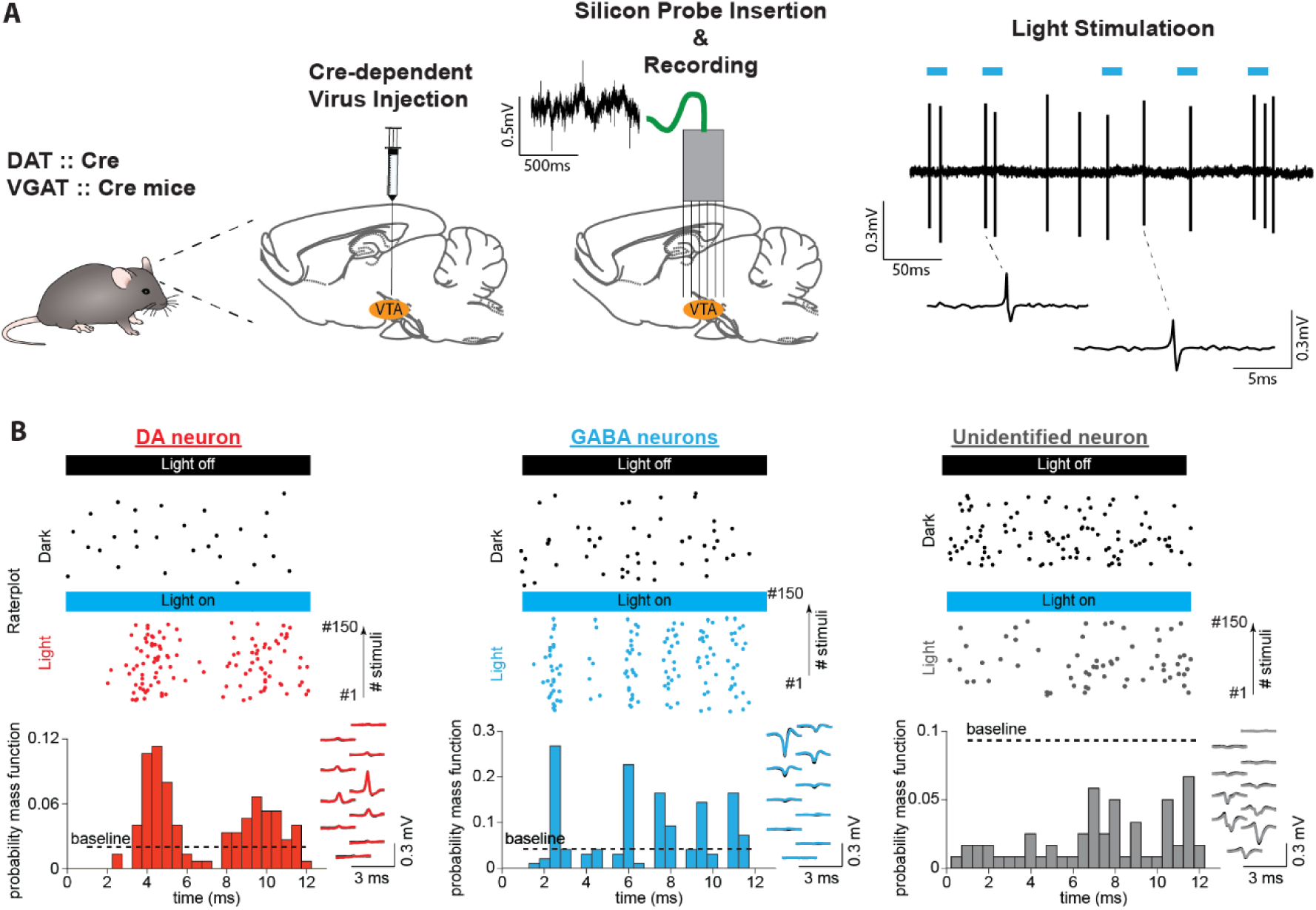
Identifying midbrain dopaminergic and GABAergic neurons. **(A)** Left: We confined ChR2 expression to DA and GABA neurons by injecting locally into the VTA the adeno-associated virus FLEX-ChR2 into transgenic mice expressing the Cre recombinase under the control of the promoter of the DA transporter (DAT::Cre) or the vesicular GABA transporter (VGAT::Cre). Approximately 10 days after the virus injection, the silicon probe was inserted into the brain in the same AP and ML coordinates. On a daily basis, the probe was inserted deeper into the brain by a few microns. Therefore, recording sessions were performed on different DV coordinates. Right: High-pass filtered voltage trace recorded during a light-stimulation session. Thick blue lines indicate light pulses (450 nm, 12 ms). Two light-induced spikes are shown below. **(B)** Light response patterns of representative DA (red), GABA (blue), and unidentified (gray) neurons. (Top) Raster plots of spikes discharged during light stimulation (colored dots) and in the inter-stimulus baseline period (baseline, black dots). (Bottom) PSTHs extracted from the light-induced spikes. The black dashed line indicates the upper confidence limit of the baseline activity. If it is exceeded by the light-induced PSTH, then the unit is identified as light-responsive (See Figure S2 for an explanation of this term). Right inset shows, superimposed, the mean waveforms of spontaneous (black) and light-induced (colored) spikes recorded by a single probe shank.

### Behavioural Performance in a memory-dependent decision-making task

Mice were trained to perform sensory-guided and memory-dependent decisions in the “Memory Task” (Figures 2A, S3A and S3B). This task required animals to associate a visual cue presented at the beginning of the trial with a rewarded side arm of a figure-eight T-maze. A short memory delay was introduced between cue presentation and action selection. Following a correct response, they received water (5 μl) from a waterspout located at the end of each arm. Depending on the individual features of cognitive demand, the maze apparatus was divided into separate sections (i.e., “start,” “cue,” “delay,” “side arms,” and “reward”). To ensure that the mice made choices guided by the visual cues and had minimal influence from other behavioral parameters on decisions, we eliminated imbalances between the left and right trials in key task parameters (e.g., reward amount, visual environment, effort, and motor skill requirements).

**Figure 2.**
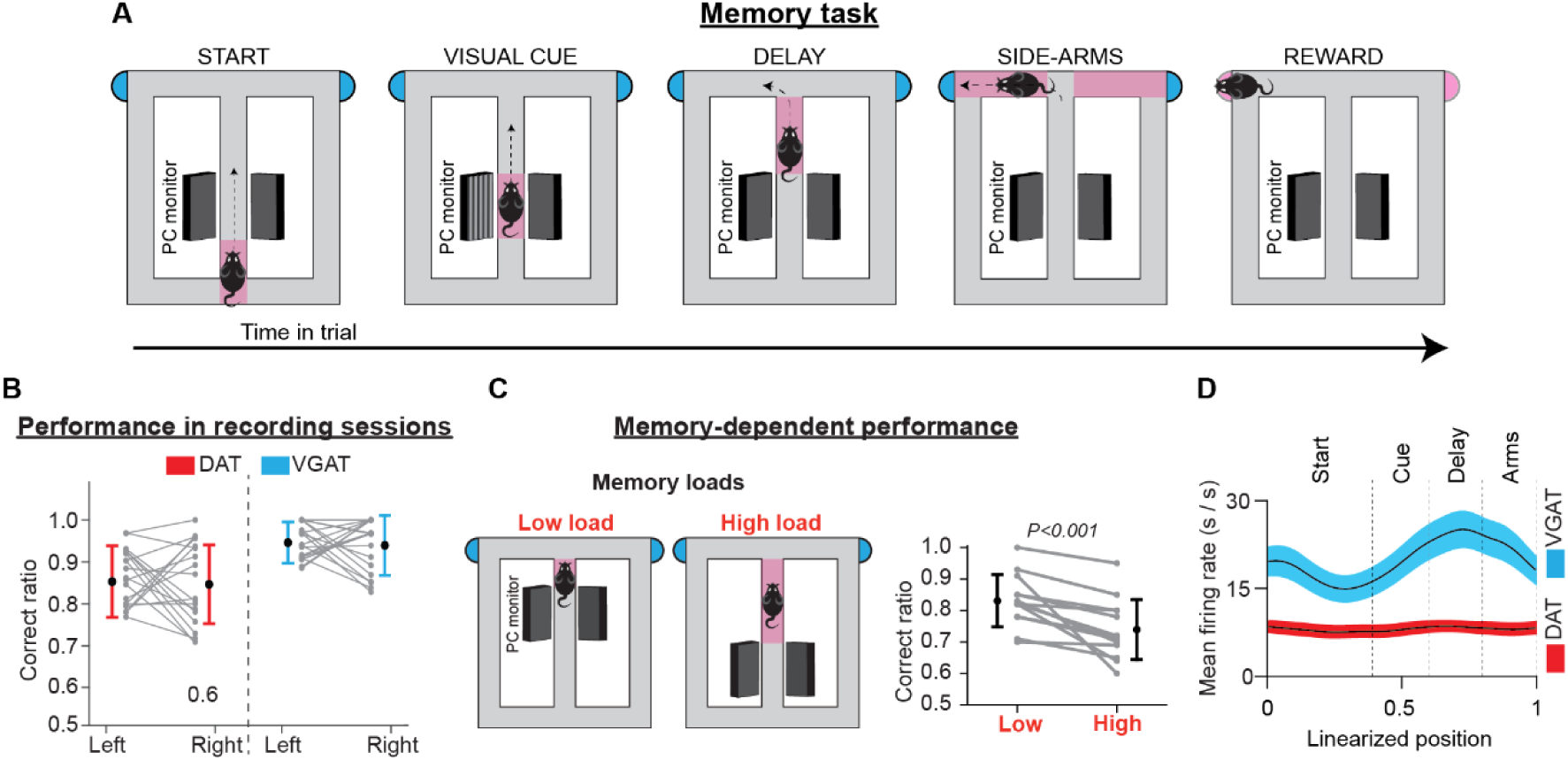
Memory task, performance, and population activity. **(A)** Schematic representation of the T-maze apparatus configuration in the memory task. Depending on the individual features of cognitive demand, the maze was divided into sections. Every trial commenced when the animal left the starting point, running along the central arm. In the visual cue section, a visual cue signaled the rewarded location. Between the visual cue section and the turning point at the end of the main arm, a brief memory delay was introduced. After reward consumption, the animals returned of their free will to the starting point to commence a new trial. **(B)** Correct performance rates in sessions with electrophysiological recordings. Gray lines illustrate performance rates for left and right trials in every session. Colored lines illustrate performance averages across sessions (mean ± standard deviation) for DAT-Cre (red) and VGAT-Cre (blue) animals. **(C)** In some training sessions animals received two blocks of trials with different memory loads. Gray lines illustrate correct performance rates for each block in every session and black lines show the average performance rates (mean ± standard deviation) across sessions, for all animals. **(D)** Mean firing rates (thick lines) ± 1 standard error of the mean (shaded areas) of the population activities of DA (red) and GABA (blue) neurons. The averaged population firing activity of GABA neurons increased in the cue and delay sections. However, the averaged population activity of DA neurons did not deviate from the beginning until the end of the trials.

At the time of neurophysiological data collection, all mice performed memory task trials with high accuracy. Averaging across sessions, the total correct rate was 86.8 ± 7.9% (mean ± standard deviation [SD]; left: 88.1 ± 10.0%, right: 87.2 ± 11.6%; paired *t*-test evaluating left vs right performance rate: t(59) = 0.46, *P* = 0.65, 60 sessions in nine mice, Figure 2B). In addition, performance was independent of individual preference for the left-or-right arm visits in any of the recorded sessions (test of independence, χ^2^(1) < 3.84, *P* > 0.05, Ho: correct rate is independent of arm choice, Figure S3C).

We also assessed the contribution of memory-related processing to task performance. To achieve this, we delivered blocks of trials with different memory loads in separate training sessions. Across all sessions, the correct performance rate dropped with higher memory load demands (mean ± SD; low load: 83.1 ± 8.3%, high load: 73.9 ± 9.5%; paired *t*-test on correct performance rate: t(12) = 4.33, *P* < 0.001, 13 sessions in seven mice, Figure 2C). This result is consistent with earlier reports (Floresco and Phillips, 2001; Floresco and Magyar, 2006) and highlights the important role of memory in supporting decisions in the present task.

### The population activity of DA neurons is not elevated during the memory task trials

DA neurons are not known to be active in the delay period of short-term memory tasks (Schultz et al., 1993; Phillips et al., 2004; Matsumoto and Takada, 2013; Choi et al., 2020), even though DA is a key neurotransmitter in the regulation of prefrontal cortical mnemonic functions (Goldman-Rakic et al., 1989; Smiley et al., 1992; Smiley and Goldman-Rakic, 1993; Goldman-Rakic, 1997; Tzschentke, 2001). This well-established notion has been established from either analysis of putative neuronal activities or inferred from voltammetry and fiber photometry recordings (Ljungberg et al., 1991; Schultz, 2002; Phillips et al., 2004; Matsumoto and Takada, 2013; Choi et al., 2020). Corroborating these earlier reports, the average discharge rate of identified DA neurons in the present study remained essentially constant (Figure 2D). Simple linear regression analysis, with the neuronal firing rate as the response variable and the animal’s position on the maze as the single predictor variable, showed that from the beginning until the end of the trial (a 1.5-meter distance), the population activity of DA neurons deviated slightly by 0.17 ± 0.62 Hz (mean ± standard error of the mean [SEM], did not differ from a distribution with a mean equal to zero; one-sample *t*-test on the position coefficient, t_(103)_ = 0.275, *P* = 0.78). Notably, in the memory-delay period, the discharge rate of DA neurons declined by - 0.72 ± 2.3 Hz (mean ± SEM, one-sample *t*-test on the position coefficient, t_(103)_ = -0.31, *P* = 0.75). On the other hand, the GABA neurons elevated their discharge rate by 4.29 ± 1.10 Hz in the delay period (mean ± SEM, one-sample *t*-test on the position coefficient, t_(73)_ = 4.09, *P* < 0.001), confirming evidence from an earlier report (Cohen et al., 2012).

### DA and GABA neurons in the VTA show trajectory-specific encoding preferences in short-term memory-dependent behavior

Making interpretations of the encoding properties of single neurons from population rate averages is highly challenging in tasks with many behavioral choices, especially for functionally heterogeneous populations such as the DA neurons. To overcome this limitation, we analyzed the firing activity of single neurons, by taking into consideration two important behavioral parameters. First, the animals visited either the left or right rewarded side arms in every trial. Therefore, we grouped and averaged trial spike trains of single neurons by the corresponding lap trajectories (left or right; see also Methods and Figure S4A). Also, in the present task, significant behavioral events (including visual cue presentation, memory delay, and reward delivery) were inherently bound to fixed positions in the maze (Figure 2A, S3A, and S3B). Thus, we arranged spiking events according to the position they occurred, to get an estimate of the behavioral correlates of neuronal activity. To this end, individual trial trajectories were linearized and represented as a one-dimensional vector consisting of 100 linearly spaced points (trial start: point 0; trial reward: point 100).

Examples of discharge patterns arranged by position and trajectory are shown in Figures 3A, S6A, and S6B. These representative neurons elevate transiently their firing activity at specific positions and do so consistently across trials. When we organized the normalized mean firing rates for the preferred and non-preferred lap trajectories of each neuron (i.e., trajectories with higher and lower firing rates respectively) by the position of elevated transient activity (left and middle heatmaps in Figure 3B) we discovered that the position preference was uniform among neurons, producing a population sequential activity from the start until the end of trials. These results of the position preference of DA neurons agree with previous studies reporting activities of DA (Engelhard et al., 2019) and cortical (Fujisawa et al., 2008; Harvey et al., 2012) neurons in the T-maze apparatus.

**Figure 3.**
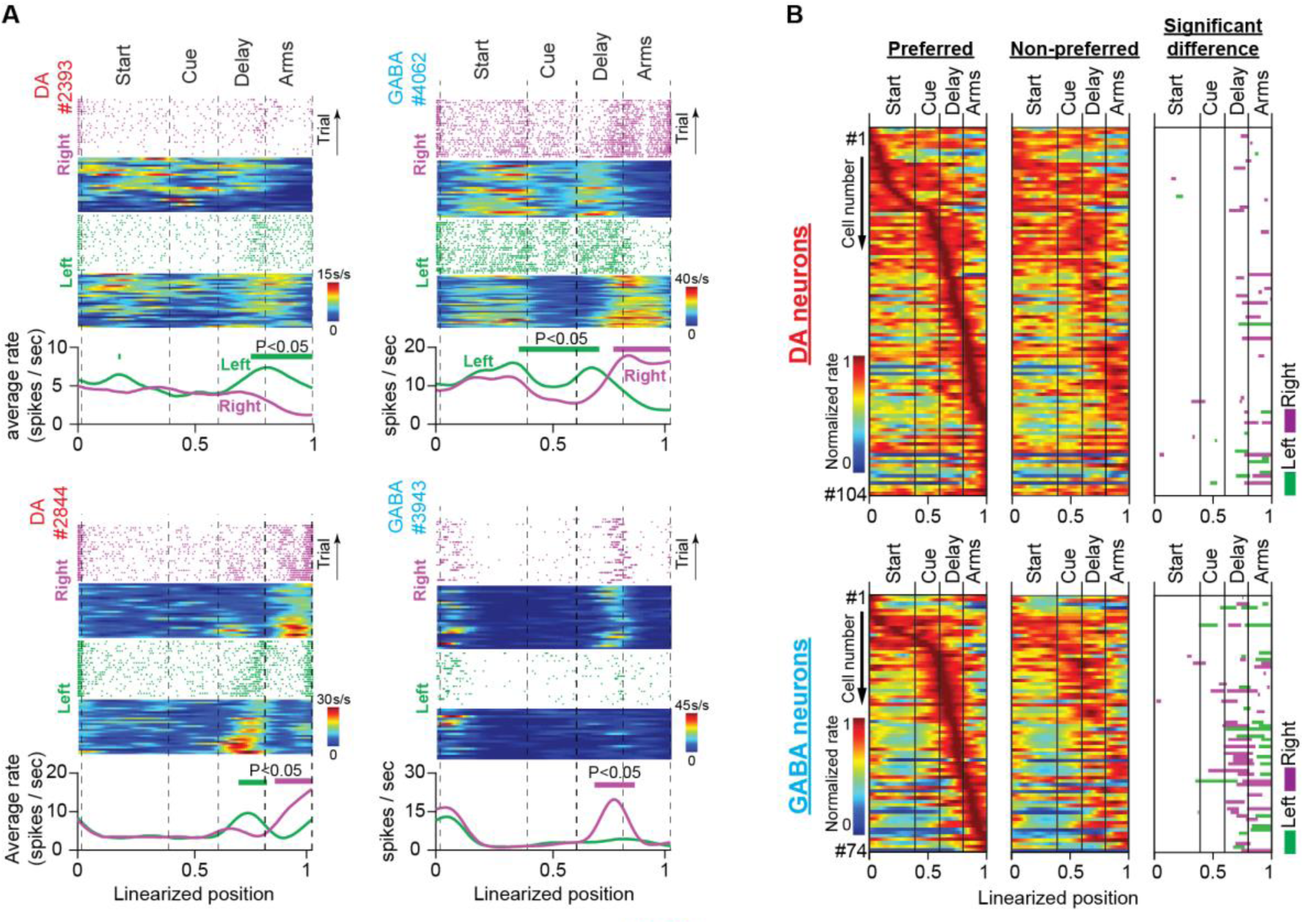
Trajectory-specific activities by DA and GABA neurons in the memory task. **(A)** Firing patterns of representative DA and GABA neurons. In each example: (Top) Raster plots of spiking events, for every correct trial, and their corresponding firing rate heatmaps as a function of position during right (purple) and left (green) trials. (Bottom) Average firing rates for correct left and right trials. Note that the trial and average firing rates (spikes/sec) are plotted as a function of position but normalized by the amount of time the mouse occupied each position on every trial. Thick lines above the average firing rates represent segments with significantly different firing rates between right and left correct trials (See also Figure S4; permutation test; *P* < 0.05). It is evident in these examples that midbrain neurons differentiate their discharge rates between left and right trajectories in certain positions. **(B)** Heatmaps of neuronal population responses organized by preferred lap trajectory (i.e., the trajectory with the stronger response; first column) and non-preferred lap trajectory (i.e., the trajectory with the weaker response; second column) for DA neurons (Top; *n* = 104 units, 35 sessions in five mice) and GABA neurons (Bottom; *n* = 74 units, 25 sessions in four mice). Each row contains preferred and non-preferred trajectory responses of the same neuron. In every row, both responses are normalized by the maximum rate of the preferred trajectory. The third column shows maze segments with significantly different discharge rates between preferred and non-preferred trajectories.

Moreover, in addition to the position preferences, a fraction of these neurons differentiated their responses between left and right trials at certain maze positions in a robust manner (Figures 3A, S6A, and S6B). To assess the trajectory-specific effects on neuronal firing activity, we used the permutation method (Figure S4). First, we calculated the original difference between the average firing rates in the left and right trials. We then randomly reassigned the trajectory labels (left or right) on the trial spike trains and produced the permuted average firing rate differences. If neurons were modulated by trajectory, the original and permuted firing rate differences were significantly different. Since spiking events were arranged by position, the permutation method could also detect positions with significant differences.

The right heatmap in Figure 3B summarizes the results from the permutation analysis applied to the populations of 104 DA and 74 GABA neurons. In both neuronal populations, there was abundant trajectory-specific activity, concentrated mostly in the delay and side-arm sections. Almost 20% of DA neurons differentiated their response between left or right trajectories in those maze sections (21% in the delay section and 22% in the side-arm section, 104 neurons, permutation test, *P* < 0.05, Figure 3B and Table S1). In GABA neurons, the percentage was even higher, with almost 50% of these cells eliciting trajectory-specific activities (47% in the delay section and 47% in the side-arm section, 74 neurons, permutation test, *P* < 0.05, Figure 3B and Table S1).

There have been reports of DA neurons discriminating between visual cues in a T-maze task (Engelhard et al., 2019) or choice selections in delayed-match-to-sample tasks (Matsumoto and Takada, 2013; Choi et al., 2020). However, in the present study, we did not detect different responses between left or right visual cues. Furthermore, neuronal preference for trajectories was not restricted to the turning point, which could indicate neuronal engagement in motor preparation for choice execution. Instead, it was spread in a wider area, covering a distance from the memory delay onset until the end of the side arms.

A plausible explanation for the trajectory-specific responses in the side arms is that neurons were under the control of the sensory, motor, or goal-directed behavioral processes triggered by the opposite trajectories (Howe et al., 2013; Hamid et al., 2016; Mohebi et al., 2019). However, in the memory-delay section, trajectories were identical for the left and right trials, which could be suggestive of the engagement of these neurons in short-term memory processing. Neuronal preferences to arm visits in memory delay are not uncommon in T-maze tasks. They have been reported in prefrontal and post-parietal cortical neurons and have been attributed to short-term memory-dependent decisions (Fujisawa et al., 2008; Harvey et al., 2012). So, is the trajectory-specific activity in our task reminiscent of internally generated, memory representations, or can be attributed to the well-known DA-linked neuronal computations (Schultz, 2002; Cohen et al., 2012; Berke, 2018; Engelhard et al., 2019)? To test this hypothesis, we proceeded to a series of statistical analyses and control behavioral tasks.

### Multiple regression analysis confirms the trajectory-specific effect on DA and GABA neurons

We discovered that significant proportions of VTA neurons fired preferentially for left or right trajectories at specific locations on the maze when we arranged discharge patterns by arm visit and position. This result does not attest that trajectory and position alone contribute to the neuronal firing rate. Midbrain DA neurons are known to respond to a wide variety of behavioral parameters (i.e., choice accuracy, reward history, running speed, and distance to rewards (Engelhard et al., 2019)) which could also exert a significant effect on neuronal firing activity. However, their effect could be dampened due to the specific firing range arrangement.

Since these behavioral variables are difficult to control with behavioral tasks, we assessed their contribution to neuronal responses using multiple regression analysis (Figure S7 and Supplementary Text 1). We found that all the examined variables (lap trajectory, trial number, speed, trial accuracy, and reward history) contributed to the firing activities of neuronal subpopulations; however, only the lap trajectory predictor could explain better the trajectory-specific activities observed in the ∼20% of DA and ∼50% of GABA neurons that were identified with the permutation analysis.

### Memory-dependent but not motivated behavior is related to trajectory-specific activity in VTA neurons

Next, we investigated the contribution of short-term memory in decision-making on the trajectory-specific activity of VTA neurons. Memory-dependent decision-making depends on three major computational components. These are (i) sensory input gating, (ii) maintaining and manipulating memory contents and (iii) generating and executing appropriate motor plans (Cohen et al., 2002; Dreher and Burnod, 2002; Montague et al., 2004; Ott and Nieder, 2019).

We eliminated all three components in a variation of the memory task. Specifically, we trained mice in the no-cue-no-choice task, in which they were not presented with a visual cue and, therefore, could not make predictions about the location of the reward (Figures 4A, S3A and S3B). Furthermore, the choice selection was prevented by the presence of blocked side arms when they arrived at the T-intersection. After a short delay (approximately 1 s), access to one of the side arms (chosen pseudo-randomly) was permitted, which always led to a reward.

**Figure 4.**
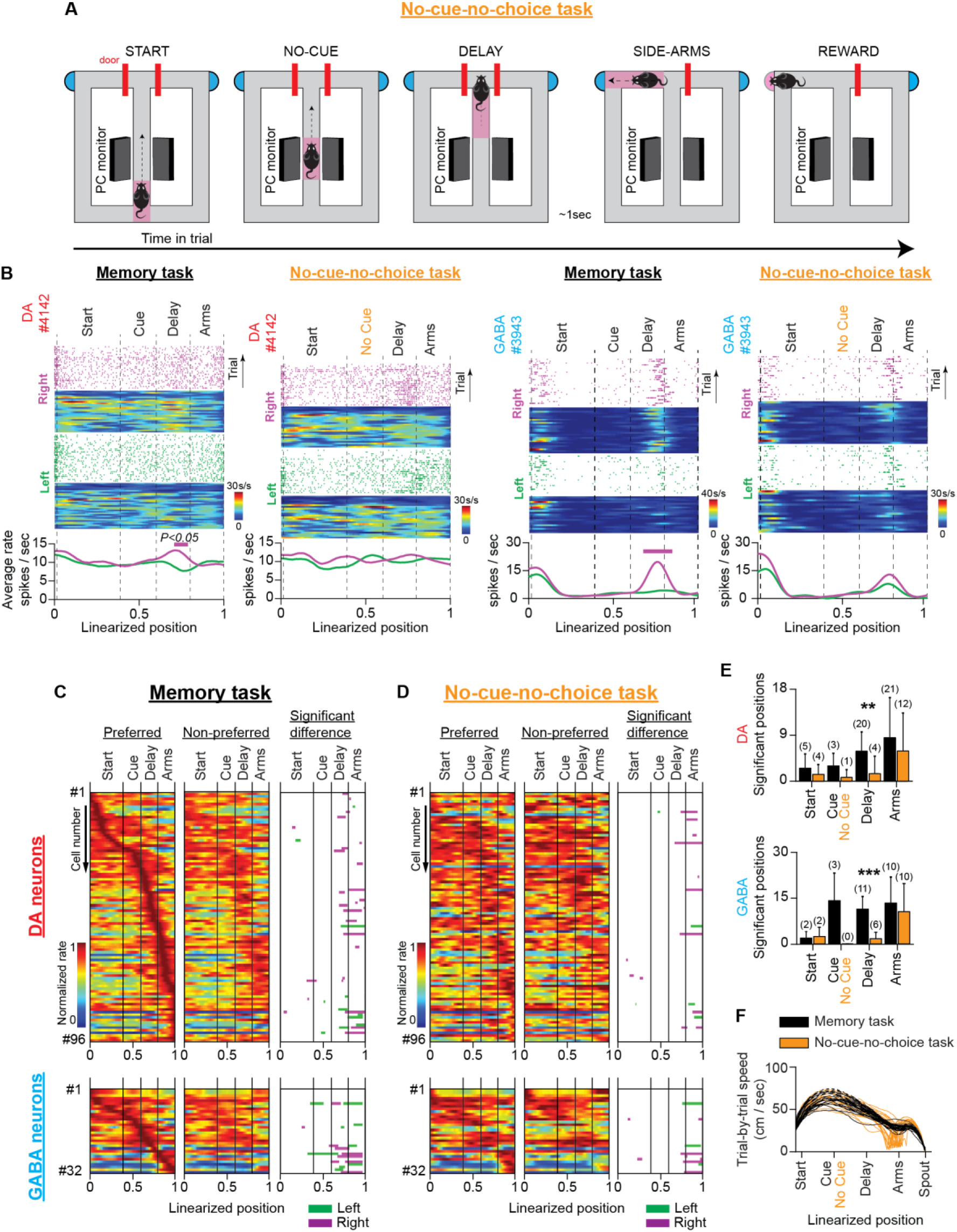
VTA neuronal responses in a T-maze task without visual cues and memory-dependent decisions (no-cue-no-choice task) **(A)** Schematic representation of the T-maze apparatus illustrating the sequence of events in the no-cue-no-choice task. **(B)** Firing patterns of representative DA (left) and GABA (right) neurons during the memory and no-cue-no-choice tasks. Both examples illustrate that the trajectory-specific firing rate difference in the delay section of the memory task declines prominently in the control task when animals do not receive visual cues which indicates the reward location, or make memory-dependent decisions. **(C and D)** The firing patterns of DA neurons (C; *n* = 96 units, 30 sessions in four mice) and GABA neurons (D; *n* = 32 units, 12 sessions in three mice) in the memory task and the no-cue-no-choice task recorded in the same sessions. (Left and Middle columns) Normalized average neuronal responses for preferred (left) and non-preferred (middle) trajectories. The right column represents the maze segments with significantly different discharge rates. The row order of the neurons is the same for the memory task and the control task heatmaps. The data shown here for the memory task are a subset of those shown in Figure 3B. **(E)** Average number of position points (mean ± standard deviation) with a significant rate difference, arranged by maze section and behavioral task for DA (left) and GABA (right) neurons (** *P* < 0.01, *** *P* < 0.001, paired *t*-test comparing numbers of significant position points between tasks. Also, the numbers in parentheses describe the number of neurons with a significant rate difference). **(F)** Representative example showing the prominent difference in running speeds (cm/s) between the memory (black) and no-cue-no-choice (brown) task trials in a single session.

In recording sessions, mice received mixed protocols composed of randomly interleaved memory task and no-cue-no-choice task trials. We evaluated the trajectory-specific activities on each task separately using the permutation method. In the delay section of the no-cue-no-choice task, we observed a significant reduction in the number of positions with a significant firing rate difference (mean ± SD; DA: memory task 5.9 ± 3.8 points, no-cue-no-choice task 1.5 ± 3.4 points, paired *t*-test, *P* = 0.002, four animals; GABA: memory task 10.7 ± 3.9 points, no-cue-no-choice task 1.5 ± 2.0 points, paired *t*-test, *P <* 0.001, three animals, Figures 4B to 4E). The attenuating effect on trajectory-specific activity was also reflected by a marked reduction in the number of trajectory-specific neurons (Figure 4E, numbers in parentheses). However, the firing rate difference between the left and right-side arms was strong in both tasks (DA: memory task 8.5 ± 7.8 points, no-cue-no-choice task 5.9 ± 7.4 points, paired *t*-test, *P* = 0.139, four animals; GABA: memory task 12.6 ± 8.1 points, no-cue-no-choice task 9.9 ± 8.8 points, paired *t*-test, *P* = 0.234, three animals, Figures 4B to 4E).

However, the significant reduction in trajectory-specific encoding preference in the no-cue-no-choice task could not be entirely attributed to the absence of the memory component. This is because important running speed, motor responses, and motivational discrepancies exist between memory and no-cue-no-choice tasks. With regard to motivation, the important role of DA in adaptive decision-making is widely recognized (Hamid et al., 2016; Berke, 2018; Mohebi et al., 2019). We did not observe animal choice bias in memory task performance (Figures 2B and S3C), but we cannot rule out the possibility that individual neurons were modulated differently by effortful actions to reach the left- and right-sided rewards (Figure S8). Unlike the memory task, in the no-cue-no-choice task, the mice could not direct behavior towards the left- or right-side arms due to the absence of a visual cue. As a result, they were unable to allocate incentive motivational drives to the left-or-right trials (Howe et al., 2013; Hamid et al., 2016; Berke, 2018; Mohebi et al., 2019). Also, the initial access denial to the side arms in the control task eliminates any potential differences in the motor preparation coding schemes (according to the “gating theory”) for the opposite arm visits in the memory task (Engelhard et al., 2019; Ott and Nieder, 2019). Finally, regarding speed, a representative example of running speed differences between the two tasks within a single session is shown in Figure 4F.

To dissociate the short-term memory component of neuronal activity from the modulatory effects of running speed, incentive motivation, and motor-related signaling, we trained mice in a second control task. The cue-no-choice task preserved the same running speed parameters (Figure S9), motor skill requirements, and physical effort demands (i.e., visual cues, maze shape, arm length, and reward amount were the same) as the memory task, but it prevented animals from making decisions. Accordingly, the animals were presented with the same visual cue as in the memory task, which indicated the side arm that was rewarded and enabled them to allocate incentive motivational drive to the left-or-right trials; however, they were always forced to visit the rewarded arm by blocked access to the unrewarded arm (Figures 5A, S3A and S3B). In the same recording session, the mice performed a separate block of memory task trials. Similar to the first control task, in the delay section of the cue-no-choice task we observed a significant reduction in the spatial extent of the firing rate difference (DA: memory task 5.4 ± 3.6 points, cue-no-choice task: 0.4 ± 1.1, paired *t*-test, *P* = 0.011, four animals; GABA: memory task 10.7 ± 6.3 points, cue-no-choice task 3.5 ± 4.2 points, paired *t*-test, *P* < 0.001, 1 animal, Figures 5B to 5E). In the side arms, however, the trajectory-specific effect remained strong and was not significantly different from the effect observed in the memory task (DA: memory task 8.1 ± 10.3 points, cue-no-choice task 5.0 ± 4.6 points, paired *t*-test, *P* = 0.146, four animals; GABA: memory task 8.7 ± 5.9 points, cue-no-choice 5.2 ± 6.2 points, paired *t*-test, *P* = 0.086, 1 animal, Figures 5B to 5E).

**Figure 5.**
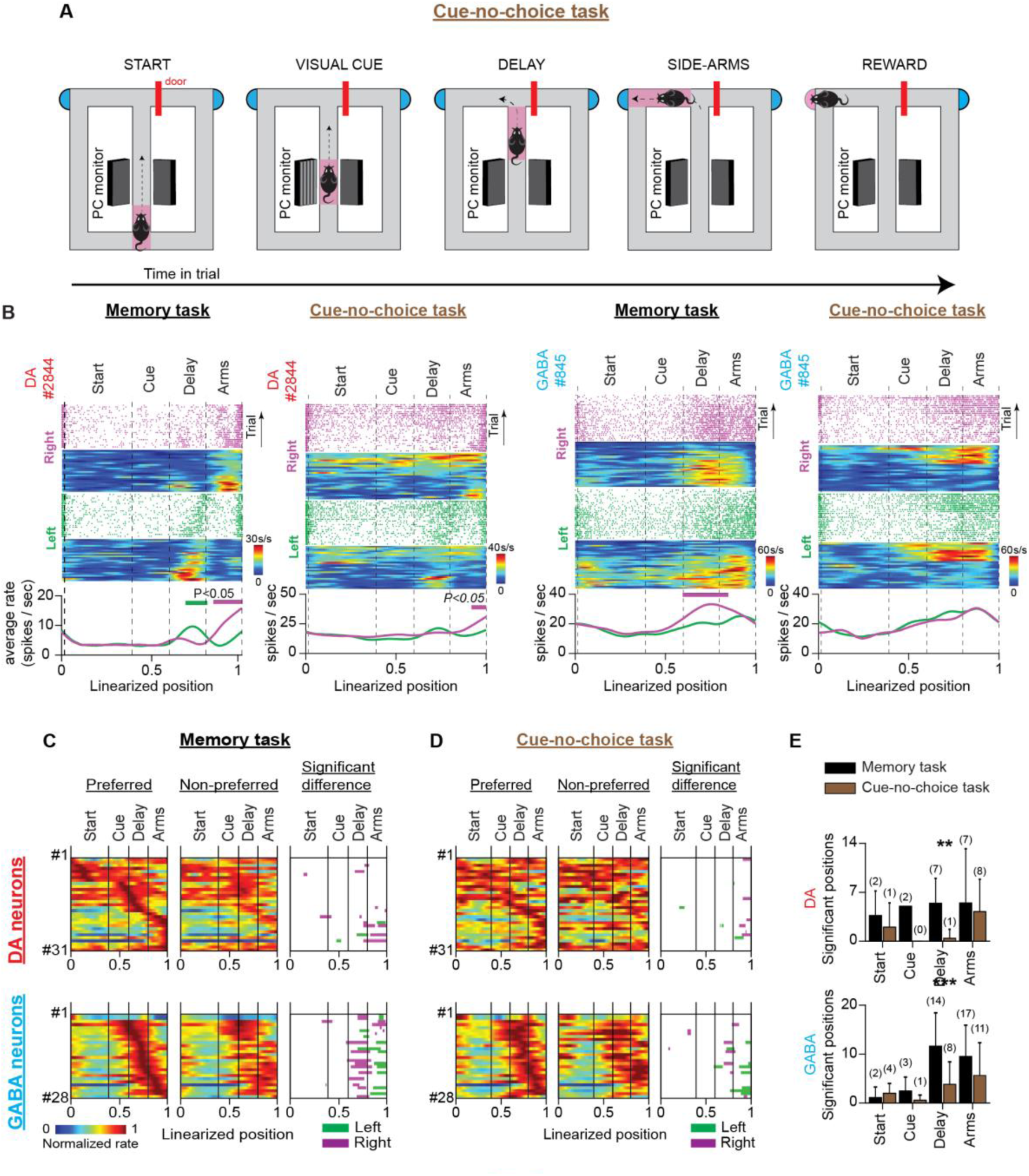
VTA neuronal responses in a T-maze task with visual cues but no memory-dependent decisions (cue-no-choice task) **(A)** Schematic representation of the T-maze apparatus illustrating the sequence of events in the cue-no-choice task. **(B)** Firing patterns of representative DA (left) and GABA (right) neurons during the memory and cue-no-choice tasks. Both examples illustrate that the trajectory-specific firing rate difference in the delay section of the memory task becomes notably weaker in the cue-no-choice task when animals do not make memory-dependent decisions, although running speed activities and incentive motivational drives of physical effort are the same between tasks. **(C and D)** The firing patterns of DA neurons (C; *n* = 31 units, 28 sessions in four mice) and GABA neurons (D; *n* = 28 units, 11 sessions in one mouse) in the memory task and the cue-no-choice task recorded during the same sessions. (Left and Middle columns) Normalized average neuronal firing rates associated with the preferred (left) and non-preferred (middle) trajectories. The right column represents the maze segments with significantly different discharge rates. The row order of the neurons is the same for the memory task and the control task heatmaps. The data shown here for the memory task are a subset of those shown in Figure 3B. **(E)** Average number of position points (mean ± standard deviation) with significant rate differences, arranged by maze section and behavioral task for (left) DA and (right) GABA neurons (** *P* < 0.01, *** *P* < 0.001, paired *t*-test comparing numbers of significant position points between tasks. Also, the numbers in parentheses describe the number of neurons with a significant rate difference).

Together, these results suggest that trajectory-specific responses in the delay period of the memory task could reflect short-term memory representations linked to decision-making behavior and cannot be explained by running speed, motor, and motivation-related signaling differences.

### Neuronal activities in delay and reward are unrelated

DA neurons are known to be excited by rewards (Schultz et al., 1993; Schultz et al., 1997; Cohen et al., 2012; Matsumoto and Takada, 2013; Engelhard et al., 2019; Choi et al., 2020). In agreement with this notion, we discovered that 27 DA neurons (28% of 104 neurons, Figure 6B column 4; we defined the first second of reward consumption as the reward section.) responded to reward with significant excitation.

**Figure 6.**
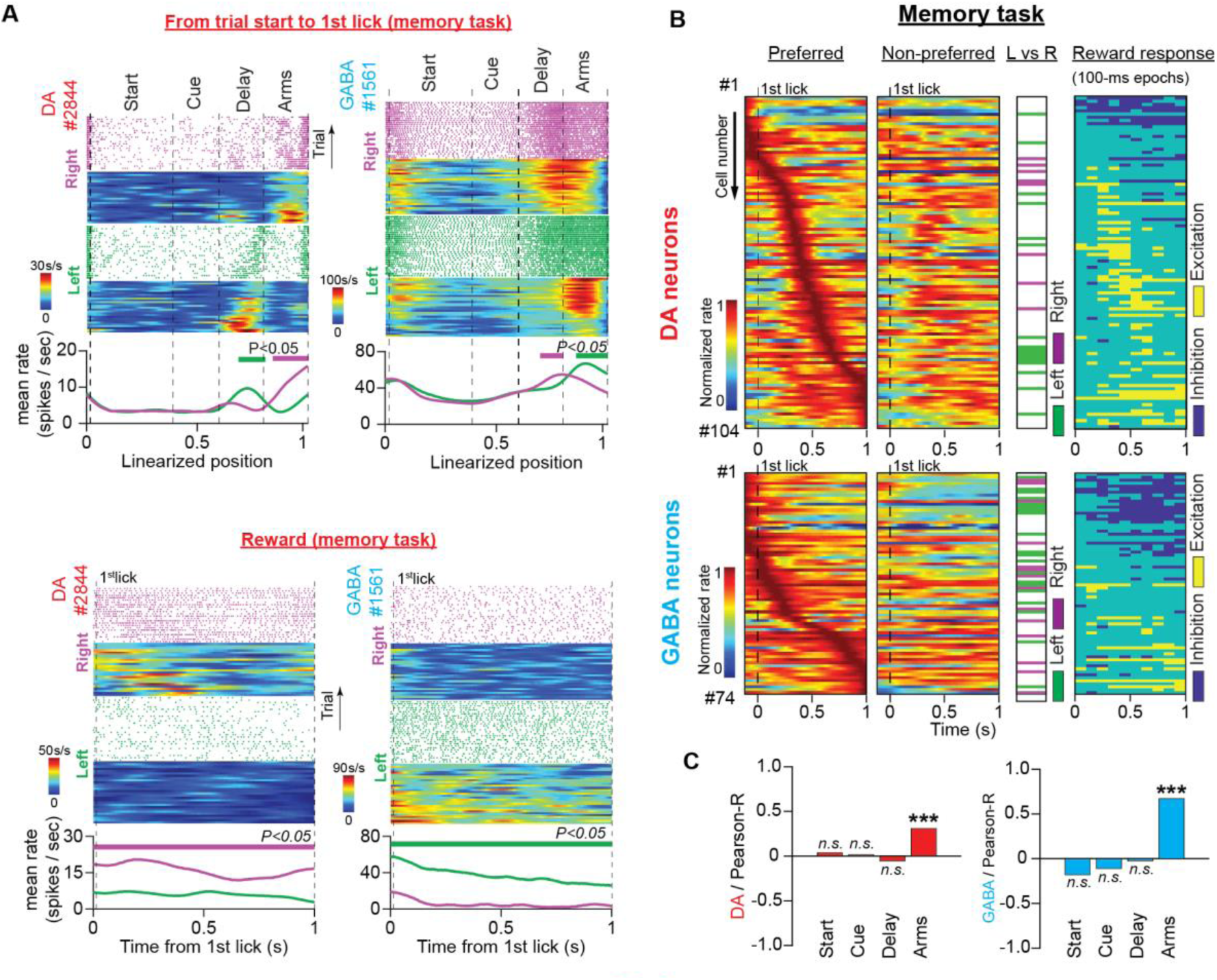
Neuronal responses during reward consumption are not related to the trajectory-specific activities in the memory delay. **(A)** Firing patterns of representative DA and GABA neurons in the memory task for the period from trial start until the first lick of the waterspout, which triggered the water pump (top, space domain) and during reward consumption (bottom, time domain). In each example: (Top) Raster plots of the spike trains and their corresponding firing rate heatmaps arranged by trajectory and position (maze) or time (reward) in right (purple) and left (green) trials. (Bottom) Average firing rates for correct left and right trials. The thick lines above the firing rates represent segments with significantly different firing rates between the correct right and left trials. **(B)** Firing patterns of DA (top) and GABA (bottom) neurons in the time domain during reward consumption (from the first lick until 1 s later) for preferred (first column) and non-preferred (second column) rewards (DA: *n* = 104 units, 35 sessions in five mice; GABA: *n* = 74 units, 25 sessions in four mice). Each row represents the normalized average firing rates (preferred and non-preferred) of a single neuron on a color scale. Neurons were ordered according to the time point of the maximum rate in the preferred arm. The third column shows neurons with significant discrepancies between the left and right reward-related responses (paired *t*-test for mean firing rates, *P* < 0.05). The fourth column shows post-delivery reward segments (100 ms each) with significant excitation or inhibition compared with the 100-ms pre-reward segment (paired *t*-test comparing firing rates, *P* < 0.05). **(C)** Correlations between the mean firing rate difference in the reward section and the difference in every other maze section for DA (left) and GABA (right) neurons (Pearson’s R values with *P*-values, *** *P* < 0.001). Only the trajectory-specific firing rate difference in the side arms correlated with the reward-specific rate difference.

DA neurons are known to discriminate between rewards with different magnitudes and predictabilities (Tobler et al., 2005; Morris et al., 2006; Matsumoto and Takada, 2013). In the present study, the animals were offered equivalent options in terms of reward magnitude, uncertainty, and effort. Thus, we predicted the presence of a small number of reward-discriminating neurons. However, we found that 23% of DA neurons and 46% of GABA neurons differed significantly in their responses to left-or-right rewards (Figures 6A, 6B column 3, and Table S1; paired *t*-test comparing mean firing rates, *P* < 0.05). This unexpected result raised the hypothesis that the trajectory-specific activities we observed in the memory delay were related to selective preference for rewards.

To test this theory, first, we correlated the average firing rate difference in the reward section with the average rate difference in the preceding maze sections. In both neuronal populations, we found a significant positive relationship between the reward and side-arm sections (Figure 6C; Pearson’s correlation; DA: *R* = 0.31, *P* < 0.001; GABA: *R* = 0.67, *P* < 0.001) but not between the reward and delay sections (Figure 6C; Pearson’s correlation; DA: *R* = -0.05, *P* > 0.05; GABA: *R* = -0.03, *P* > 0.05).

Next, we sought to determine whether neurons with trajectory-specific activities in memory delay, also exhibited a significant preference for the same-trajectory reward. To do so, we divided neuronal encoding preferences in six (6) categories determined by the maze section they elicited significant firing differences (delay or reward) and by the preference for trajectory (left-significant, right-significant, or non-significant). We discovered that only four (n= 4 out of 104) DA neurons showed the same side preference in the reward and delay sections (e.g. elicit significantly stronger firing activities in the delay section of left trials and left reward). Thirty-one (n=31) neurons responded differently and sixty-nine (n=69) did not elicit significant responses in any of the sections (Figure S10).

The results from both analyses converge to the conclusion that trajectory-specific firing activities in memory delay do not reflect reward preference during consumption.

### Anatomical organization of trajectory-specific neurons

Several recent studies have reported that neighboring DA neurons are more likely to share similar encoding properties, thus, forming functional but also anatomical clusters within the VTA (Lammel et al., 2008; Matsumoto and Takada, 2013; Engelhard et al., 2019). Therefore, we sought to determine whether neurons with memory-specific encoding properties were anatomically segregated from the rest of the population. Estimating the location of recorded neurons (i.e. Bregma vs Mediolateral coordinates) from the anatomical reconstruction of the recording channels revealed that the trajectory-specific GABA neurons were located more laterally compared to the rest of the group (Bregma: t_(72)_ = 1.165, *P* = 0.248, Mediolateral: t_(72)_ = -2.38, *P* = 0.019, Figure S11). However, in the DA neurons there was no clear anatomical segregation (Bregma: t_(102)_ = 0.045, *P* = 0.964, Mediolateral: t_(102)_ = 0.177, *P* = 0.860, Figure S11). The lack of evidence of functional and anatomical segregation in the DA neurons could be accounted for by the fact that we targeted mostly the lateral parts of the VTA. Also, since the position of the recording channels along the dorsoventral axis was changing on a daily basis, we did not include in our analysis the dorsoventral coordinates of the recorded neurons.

## Discussion

In the present study, we performed extracellular recordings from optogenetically identified DA and GABA neurons in the VTA while mice performed reward-seeking tasks on a T-maze apparatus. Mice were trained to choose between two spatially separate goals under the instruction of visual cues presented at the beginning of the trial. A short memory delay was introduced between cue presentation and choice selection. We discovered that subpopulations of DA and GABA neurons showed differential responses between the left and right trials, starting from the onset of the memory delay in the main arm, where the trajectories were indistinguishable. Trajectory-specific preference was not correlated with reward history, running speed, the incentive motivational drive of physical effort, or reward-related encoding differences, and diminished significantly when the memory-dependent decision component was eliminated in control behavioral tasks. This evidence indicates that populations of DA and GABA neurons in the VTA encode internally generated signals that support short-term memory in decision-making.

### Activities of midbrain DA neurons in short-term memory

The “gating theory” unifies the signaling activities of DA neurons in reward prediction and short-term memory (Cohen et al., 2002; Dreher and Burnod, 2002; Montague et al., 2004; Ott and Nieder, 2019). With regards to mnemonic processing, the well-established notion that DA somatic spiking activity is low in short-term memory stemmed either from recordings of putative DA neurons of the A8, A9, and A10 pathways (Schultz et al., 1993; Schultz, 2002; Matsumoto and Takada, 2013) or inferred from neuronal population activities (Phillips et al., 2004; Choi et al., 2020). Consistent with the latter reports we did not observe a profound variation in the population activity of DA neurons during the memory task.

However, a wealth of recent studies has shown that DA neurons are functionally and genetically segregated (Lammel et al., 2008; Lammel et al., 2011; Engelhard et al., 2019). Moreover, in many real-life situations, animals must choose between many options for behavioral responses in high-dimensional environments. In such behavioral conditions, averaging neuronal responses, irrespective of the behavioral features and decisions they respond to, could hinder the fine computational processes of single neurons. To overcome this limitation, we analyzed the firing activities of identified single neurons, focusing on different discharge patterns between behavioral choices. Here, we demonstrated that memory-specific activities by midbrain DA neurons can be represented as trajectory-specific responses in the delay period of the memory task. Other than gating sensory input, and maintaining and manipulating memory contents, DA has been implicated in relaying motor commands to elicit memory-guided responses (Matsumoto and Takada, 2013; Engelhard et al., 2019; Ott and Nieder, 2019). We tested this theory, but we found that the firing rate differences between left and right-arm responses declined in a control behavioral task (cue-no-choice) without memory load but with the same motor skill requirements as the memory task. From this, we concluded that motor preparation coding schemes for left or right arm responses (Cohen et al., 2002; Ott and Nieder, 2019) cannot account for the trajectory-specific activities in the memory delay of the T-maze task. Instead, our evidence indicates that trajectory-specific activities by DA (also GABA) neurons are functionally linked to the maintaining and manipulating of memory contents.

Our study provides more evidence to a mounting body of recent work which suggests a dynamic functional interaction between the PFC and VTA circuits in high dimensional behavioral environments. The memory-related, trajectory-specific midbrain neuronal activities demonstrated here, have also been reported for PFC and post-parietal neurons while rodents perform reward-seeking responses on the T-maze (Fujisawa et al., 2008; Harvey et al., 2012). We also report that while mice navigate the maze, individual VTA neurons elevate transiently their firing activity at unique positions producing at the population level a neuronal sequential activity, which is a well-known physiological hallmark of cortical neurons. PFC and VTA networks are known to oscillate in high synchrony at 4 Hz in T-maze tasks (Fujisawa and Buzsáki, 2011). Network oscillations were also evident in our VTA recordings (unpublished data). Finally, we report that DA and GABA neurons of the VTA exhibit multitasking activities, encoding behaviors that overlap those of PFC neurons (see further discussion below). These striking similarities are in line with a recent computational theory proposing that the encoding properties of DA neurons reflect those of the upstream PFC neurons (Lee et al., 2022).

The present study also corroborates important findings from a recent report, which demonstrated that optogenetic perturbations in DA neuron excitability exert a strong effect on short-term memory performance, highlighting the causal role of DA neuronal firing activity in memory-dependent behavior (Choi et al., 2020). Also, in agreement with a previous report (Engelhard et al., 2019) by the same laboratory, we show that subpopulations of VTA neurons are modulated by running speed, cumulative performance rate, current choice accuracy, and reward history. Disparities between this and our study in the proportions of modulated neurons could be attributed to the different recording techniques applied as well as the maze regions of interest. Although, Engelhard et al. trained mice in a virtual T-maze task, they analyzed neuronal firing activities and identified choice-specific neurons only in the visual-cue period, but not in memory delay (Engelhard et al., 2019). In contrast, we focused our analysis on the memory delay of the T-maze task.

Overall, our results agree with the notion that DA neurons encode a variety of behavioral parameters in complex environments. In addition, we confirmed that in memory-dependent behaviors, DA neuronal populations did not elicit sustained increases in their discharge rate. However, in the present task, DA neurons individually encoded internal representations by differentiating their responses to lap trajectories in memory delay.

### Role of motivated behavior in trajectory-specific encoding properties of VTA neurons

Midbrain DA activity is known to be involved in motivated behavior while rodents navigate mazes to receive rewards beyond immediate reach (Hamid et al., 2016; Berke, 2018; Mohebi et al., 2019). When mice approach rewards, striatal DA concentrations increase, scaling flexibly with reward size and proximity, which is proposed to reflect a neural correlate of a sustained motivational drive (Howe et al., 2013). To evaluate the role of motivated behavior in the trajectory-specific preference of midbrain neurons, we compared firing activities between a memory task and a control task without memory-dependent decisions (cue-no-choice task). Although in the cue-no-choice task, the behavioral parameters that determined the incentive motivational drives were the same as in the memory task (visual cues, maze shape, and reward amount), neuronal responses did not differ between the left and right trials. This result strongly indicates that incentive motivational drives (at least for physical effort) do not contribute to trajectory-specific activities of midbrain neurons during the delay period of the memory task.

### Memory-specific activities of the VTA neurons are not attributed to reward prediction error signaling

We also assessed the role of reward-related processing in the trajectory-specific activity of the midbrain neurons. When animals estimate the spatial proximity of distant rewards, DA neurons calculate RPE signals from state-value functions (Hamid et al., 2016; Berke, 2018; Engelhard et al., 2019; Mohebi et al., 2019; Kim et al., 2020). In the present study, the animals received ongoing visual input, facilitating the continuous estimation of reward proximity. Thus, DA neurons can potentially estimate RPE signals from successive state values assigned to each position on the maze track (Figure S8). Therefore, the difference in firing activity between the left and right trials could be the result of differences in the state-value functions assigned to these trajectories (Hamid et al., 2016; Berke, 2018). However, significant evidence contradicts this hypothesis.

First, the behavioral parameters that determine the state-value functions for the left and right trajectories were set to be identical in the cue-no-choice task and memory task, by preserving the same maze configurations and delivering equal amounts of reward. In addition, behavior in both tasks was cue-driven; therefore, animals could make predictions about the reward location and orchestrate behavior accordingly. However, we observed a prominent reduction in the firing rate difference between the left and right trials in the cue-no-choice task (Figure 5). Second, a significant subset of DA neurons (approximately 20%) responded differently to the left and right rewards in the memory task, although the same amount of reward was delivered. This unexpected finding raised the hypothesis that the encoding preference for reward could be reflected in the values of the preceding states in the maze and, therefore, could account for the trajectory-specific effect in memory delay. However, the differences in the firing activity elicited by the consumption of left-or-right rewards were unrelated to the firing rate difference in the delay section (Figure 6C). Also, we discovered a very small minority of neurons with the same trajectory preference in the memory delay and reward sections within the same trial (Figure S10B). In conclusion, these findings indicate that the encoding preference for lap trajectories exhibited by midbrain DA and GABA neurons cannot be simply explained by discrepancies in RPE signaling.

### GABA neurons of the VTA and short-term memory

With the advent of highly selective identification and perturbation techniques, new evidence has emerged regarding the encoding properties and functional roles of local VTA inhibitory networks in reward processing and motivation. There are reports demonstrating that GABA neurons of the VTA suppress reward consummatory behavior (van Zessen et al., 2012), facilitate aversive behavior (Matsumoto and Hikosaka, 2007; Tan et al., 2012), and elicit sustained activities in the delay period between conditioned and unconditioned stimuli (Cohen et al., 2012). During these behaviors, the responses of DA and GABA neurons are often inverse, such that when GABA neurons are excited, neighboring DA neurons decrease their discharge rate. In particular, aversive stimuli excite GABA neurons, which then suppress the neighboring DA neurons (Tan et al., 2012). In addition, during reward consumption, GABA neurons are inactive (Cohen et al., 2012; van Zessen et al., 2012); however, when excited, they inhibit DA neurons and disturb consummatory behavior (van Zessen et al., 2012). Finally, in classical conditioning tasks, DA neurons respond to rewards and reward-predicting stimuli, whereas GABA neurons remain silent during such events (Cohen et al., 2012). However, we demonstrated here that midbrain DA and GABA neurons elicit remarkably similar encoding properties. Both neuronal populations respond to short-term memory-specific activities manifested by encoding preferences for lap trajectories. Notably though, GABA neurons are more strongly engaged in this dynamic encoding activity since almost twice as many inhibitory neurons responded differently to the left and right trials.

This result presents an activity paradox. Given the abundant and potent synaptic inhibition of DA neurons by neighboring GABA neurons (Omelchenko and Sesack, 2009), it was unexpected that both populations were highly active and similarly engaged in tasks. However, anatomical evidence provides a plausible explanation. Local inhibitory neurons form a dense network of local synaptic innervations that target the dendritic sites of DA and other GABA neurons (Traub et al., 2004; Buzsaki, 2006; Omelchenko and Sesack, 2009). Although potent and well-suited for coordinated network activity, this synaptic inhibition is not as strong as somatic inhibition (Jhou et al., 2009; Omelchenko and Sesack, 2009), and it has been suggested that it is not sufficient to suppress DA neurons when they receive a strong excitatory drive from extrinsic sources (van Zessen et al., 2012).

Finally, the stronger engagement of GABA neurons in trajectory-specific activity is an interesting observation which requires further investigation. In our opinion, future research on this topic should point to the direction of network oscillations. The VTA circuit is known to oscillate in memory-engaging behaviors producing frequencies of a wide spectrum; up to 100 Hz (Fujisawa and Buzsáki, 2011); also unpublished observations in our study). Although the mechanisms supporting circuit oscillations in the VTA are not well investigated, evidence from the prefrontal cortex (van Aerde et al., 2008; Glykos et al., 2015) and the hippocampal formation (Traub et al., 2000; Mann and Paulsen, 2007) demonstrate that the excitation of the local network of inhibitory neurons is crucial for the generation and maintenance of network oscillations.

## Conclusion

In summary, we optogenetically probed DA and GABA neurons in the VTA while mice performed a decision-making task with memory load. We discovered that both neuronal populations elicited memory-dependent preferences for left-or-right trajectories that could not be explained by motor activity, motivated behavior, or reward-related processes. This evidence indicates that VTA neurons encode mental representations to support short-term memory-dependent decisions and provides insights into novel sophisticated coding strategies employed by the midbrain DA and GABA neurons in reward-related behavior.

## Methods & Materials

### Key resources table

| REAGENT or RESOURCE | SOURCE | IDENTIFIER |
| --- | --- | --- |
| Antibodies |  |  |
| Anti-Tyrosine Hydroxylase | EMD Millipore |  |
| Anti-GFP | Aves Labs |  |
| Alexa Fluor 546 IgG | Thermo Fisher Scientific |  |
| Alexa Fluor 488 IgG | Jackson ImmunoResearch Laboratories, Inc. |  |
| DAPI Fluoromount-G | SouthernBiotech |  |
| Bacterial and virus strains |  |  |
| AAV5-EF1a-DIO-hChR2(H134R)-EYFP-WPRE-pA | UNC Vector Core | N/A |
| Experimental models: Organisms/strains |  |  |
| Mouse: C57BL/6J-Slc6a3<sup>tm1.1(cre)Bkmn/J</sup> | The Jackson Laboratory | Jax #006660; RRID: IMSR_JAX:006660 |
| Mouse: C57BL/6J- Slc32a1<sup>tm2(cre)Lowl</sup> | The Jackson Laboratory | Jax #0016962; RRID: IMSR_JAX:0016962 |
| Software and algorithms |  |  |
| MATLAB | MathWorks | https://www.mathworks.com/ |
| KlustaKwik2 | (Kadir et al., 2014) | https://github.com/kwikteam/klustakwik2 |
| RPvdsEX | Tucker-Davis Technologies | https://www.tdt.com/component/rpvdsex/ |
| Other |  |  |
| Silicon probe: 64-site, 6-shank | NeuroNexus | Buzsaki64spL |
| Optical fiber AFS50/125Y | Thorlabs, Inc. | https://www.thorlabs.co.jp/thorproduct.cfm?partnumber=AFS50/125Y |
| Blue laser diode | OSRAM Opto Semiconductors | PL450B |
| 256-channel Multiplexed Biosignal Amplifier | Amplipex, Ltd. | KJE-1001 |
| Figure-eight T-maze apparatus | O'Hara & Co., Ltd. | http://ohara-time.co.jp/ |

### CONTACT FOR REAGENT AND RESOURCE SHARING

Further information and requests for resources and reagents should be directed to and will be fulfilled by the Lead Contact, Dr. Shigeyoshi Fujisawa.

#### Animals

All experiments were approved by the RIKEN Institutional Animal Care and Use Committee. We used five adult male DAT-ires-Cre (Jackson’s Lab; stock #6660; (Bäckman et al., 2006) and four Vgat-ires-Cre (Jackson’s Lab; stock #16962; (Vong et al., 2011) mice backcrossed to C57BL/6J. Animals were housed in separate cages on a 12-h dark/light cycle and each performed the behavioral tasks at the same time of the day, between 11:00 and 17:00. In the cage, they were provided *ad libitum* food access but were restrained from water availability.

#### Behavioral tasks

All behavioral tasks took place on a T-maze apparatus. More information about the maze configuration is provided in Figure S3.

##### Memory task

Behavioral sessions commenced with the animal being placed at the “starting position” (Figures 2A, S3A and S3B). Then, access to the main corridor was provided and the animal had to run through the “start” section (0-50 cm) before it arrived at the maze segment surrounded by two PC monitors (“visual-cue” section, 50-80 cm). In this section, it was presented with a distinctive visual object (vertical black and gray bars) in one of the two monitors (the other monitor remained dark) indicates which side arm to visit to obtain the reward (i.e., left cue → left reward, right cue → right reward). In the third region of the central arm (“delay” section, 80-120 cm) both monitors turned dark. While running in the delay section, the animal had to maintain the reward-related information and based on that perform the action selection at the T-intersection. The intersection at the end of the main arm designated the end of the delay section and the beginning of the “side-arms” section (120-150 cm) where the animal runs towards the reward position in anticipation of the reward. Reward (5 μl water) was delivered on correct trials at the end of the side-arms section from a waterspout. The first activation of the light-beam sensor at the waterspout triggered the water-delivery pump (Burkert, Ingelfingen, Germany), followed by reward consumption (“reward” section). After consuming the reward, the animal could return of its own will to the starting position, to commence a new trial.

Daily behavioral sessions consisted of 80-100 trials. Only animals with at least three successive sessions with an 80% performance ratio or more in the training phase were been proceeded to surgical operations.

##### No-Cue-No-Choice task

Trials of this control task were delivered in recording sessions, interleaved with memory task trials. When the animal entered the visual-cue section it was not instructed by the visual cue (Figures 4A, S3A and S3B). Also, access to both side arms was initially denied by closed sliding doors. Approximately 1 s after the animal arrived at the turning point, one of the sliding doors opened (pseudo-randomly) providing access to the reward. On every trial, the presentation or absence of the visual cue could instruct the animal about the task rules (i.e., memory task or no-cue-no-choice task).

##### Cue-No-Choice-task

The settings of this control task were the same as the memory task settings, except for the blockade of the unrewarded side arm (Figures 5A, S3A and S3B). Thus, the animals were always forced to perform correct choices. Because in both tasks, the same cue was presented, the animals could potentially be confused about the trial’s task rules. Therefore, memory task and cue-no-choice task trials were delivered in separate sets within the same recording session. Accordingly, when the animals completed the set of cue-no-choice task trials (approximately 50 trials), they were automatically delivered with another set of memory task trials (approximately 50 trials).

Recording sessions lasted approximately 20-30 minutes.

#### Intracranial surgeries and electrophysiological recording

The surgical process consisted of two separate operations. First, mice (DAT-ires-Cre or Vgat-ires-Cre) were surgically injected with 200-500 nl of adeno-associated virus AAV5-EF1a-DIO-hChR2(H134R)-EYFP-WPRE-pA (University of North Carolina vector core facility; (Tsai et al., 2009) into the VTA stereotaxically (from inferior cerebral vein AP: ∼6.65 mm, from midline ML: - 0.55 mm on the left hemisphere, from surface 4-4.5 mm, Figure 1A). Ten to fifteen days later, mice were implanted with a silicon probe in the same AP and ML coordinates (vertical insertion was intended, 0 degrees; Figures 1A, S1A to S1D). We used Buzsaki64spL (NeuroNexus, Ann Arbor, MI, USA) silicon probes which are composed of 6 shanks (10 mm long, 15 μm thick, 200 μm shank separation) and each shank has 10 recording sites (160 μm^2^ each site 0.6- 1.0 MΩ impedance). The silicon probe was attached to a custom-made micromanipulator and moved gradually to the desired depth position. On every probe shank, optic fibers were firmly attached to secure an accurate and firm insertion of the recording channels in the deep midbrain area (Figure S1B). For experiments where light delivery was required, two of the optic fibers (shanks 2 and 5) were coupled with blue (450 nm) laser diodes (PL450B, OSRAM Opto Semiconductors). Light dispersion could potentially cover the axial and transverse span of all 64 channels (Stark et al., 2012).

During recording sessions, the wide-band neurophysiological signals were acquired continuously at 20 kHz on 256-channel Amplipex systems (KJE-1001, Amplipex Ltd, Hungary; (Berényi et al., 2014)). Following surgery, the probe was inserted 45 μm deeper into the brain daily, until it reached the VTA. Thereafter, the probe was moved deeper by 20 μm / day. The average recording coordinates for the DAT-Cre animals are 3.32 ± 0.32 mm (mean ± standard deviation) rostrocaudal and 0.82 ± 0.17 mm mediolateral, and for the VGAT-Cre animals, 3.52 ± 0.29 mm rostrocaudal and 0.90 ± 0.17 mm mediolateral (Figure S1A).

We cannot exclude the possibility that some neurons were recorded in successive sessions because clustering analysis was performed on individual sessions.

#### Data analysis

Unless otherwise stated, data analysis was performed with custom-made programs designed in MATLAB with Signal-processing and Statistics toolboxes.

##### Light-stimulation protocols for optogenetic identification

Light stimulation protocols were delivered before and after the behavioral tasks. They were composed of 1, 2, 3, and 4 mW blocks of 450 nm light pulses. Each block consisted of 150 square pulses (12 ms pulse duration; 0-1 ms and 11-12 ms contained artifacts) delivered at 1, 2, 3 to 10 Hz. Electrophysiological data recorded during light stimulation and behavioral protocols within a single session were merged and clustered together.

##### Statistical analysis for detection of light-responsive units

Neurons with light-induced responses exceeding the average spontaneous activity were classed as light-responsive. To identify light-responsive neurons we applied the statistical analysis described in detail in Figure S2.

##### Estimation of firing activity during behaviour

To estimate the neuronal firing activity while animals performed the behavioural task, we took into consideration the primary goal of this study; that is to look for trajectory-specific encoding properties, as well as the inherent limitation of the task; that is the experimenter could not control the temporal precision of the behavioural events. To overcome this limitation, we arranged firing activity by the animal’s position on the maze. To do so, first, we linearized the trial trajectories and assigned them with a lap trajectory label (left or right). Then, the linearized products were divided into 100 position points and normalized so that position 0 corresponded to the starting point of the trial and position 1 to the waterspout. Second, we constructed post-distance histograms, analogous to the peri-stimulus-time-histograms (PSTHs), although the time of spiking events was replaced by the position they occurred (for the purpose of simplicity, also by habit, we will call the post-distance histograms as PSTHs). To construct accurate PSTHs we considered the exact position the spikes were discharged and the time the animal occupied this certain position. Let *n*^(*k*)^(*x*) be the number of spikes of a single neuron and *t*^(*k*)^(*x*) be the occupation time in the *x_th_* position point of the *k_th_* trial (Figure S4A). Then, 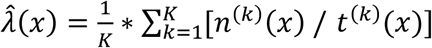 where *K* is the number of trials, represents the average firing rate probability (spikes / sec) at position point *x*. To examine the trajectory-specific encoding properties of VTA neurons we produced average firing rate histograms for correct left and right trials, separately. Then, both histograms were smoothed with a Gaussian Kernel function (σ= 0.5, length of 20 position points).

##### Firing rate heatmap construction

To construct the normalized firing rate heatmaps shown in Figures 3B, 4C, 4D, 5C, 5D, 6B, S5C to S5E and S6B we took the following steps. First, for every neuron we produced the average firing rate for left and right correct trials. Second, we normalized both rates by dividing them with the maximum firing rate of the strongest trajectory response (e.g., for the example shown in Figure S4A we divided both average firing rates by the maximum rate of the response to the left trials). Then, the normalized rate of the stronger trajectory response was assigned to the “preferred” heatmap and the rate of the weaker trajectory response to the “non-preferred” heatmap (e.g., for the example shown in Figure S4A, the left normalized rate was assigned to the “preferred” heatmap and the right rate to the “non-preferred” heatmap). Both rates occupied the same row. The row ordering was determined by the position of maximum rate.

##### Identifying trajectory-specific neurons with the permutation method

To identify neurons with trajectory-specific encoding properties we applied the permutation test reported elsewhere (Fujisawa et al., 2008) and described in detail in Figure S4.

##### Regression analysis

We designed a generalized linear regression model (GLM) with the neuronal firing rate (FR) modelled as a gaussian function of the lap trajectory (T), speed (S), trial number (TN), performance (R), current trial accuracy (A_0_) and previous trial accuracy (A_-1_) behavioural variables. With the permutation analysis we observed that the trajectory-specific effect on the firing activity was dependent on position. Thus, we examined the joint effect of trajectory with position (P) on spiking activity. All dependent and independent variables were arranged by position. The values of the trajectory (1 for left and 2 for right), trial number, performance (cumulative correct rate), current trial and previous trial accuracy (1 for correct trial, 0 for error trial) variables remained constant throughout the whole trial. The firing rate, position and speed variables changed their values on every position.

The GLM was:

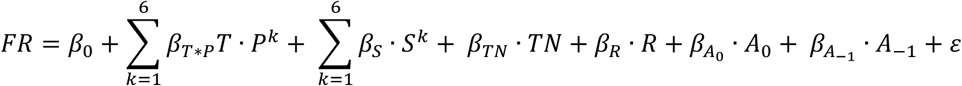

where the β values are the regressor coefficients for the different predictors (including the intercept β_0_) and ε is the Gaussian noise term. The 6^th^ degree order polynomials of position and speed were chosen for model optimization with the Bayes information criterion.

First, we generated model predictions of the average firing rates for left (*L̂*_0_) and right (*R̂*_0_) trials, and from those we calculated the predicted firing rate difference (*D̂*_0_). Then, we shuffled the trajectory labels assigned to the tested variable (the assigned labels to the rest of the independent variables remained intact) and assessed the effect on the firing rate difference. For every predictor we produced 500 shuffled rate differences, *D̂*_*j*_. If the absolute mean value of *D̂*_0_ exceeded the top 5% of the *D̂*_*j*_ values (including Bonferroni correction), then the hypothesis was rejected, and the predictor was significantly contributing to the firing rate difference. We examined every maze region individually, but here we report only for the delay region.

##### Reward-related excitation or inhibition

The reward section was defined as the first second of reward consumption. To assess neuronal response to reward consumption and categorize it as excitatory, inhibitory, or non-responsive we performed the following analysis. First, we produced the smoothed mean firing rate response in the time domain (as we did in the maze sections in the space domain) for left, ^*λ̂*^_*Left*_(*t*), and right, ^*λ̂*^_*Rig*ℎ*t*_(*t*) trials. For the preferred arm of each neuron, we compared the mean firing rate in the reward section to the mean rate in the 100 ms epoch preceding reward delivery (paired t-test on mean firing rates; *P < 0.05*; Figure 6B column 4).

##### Encoding preferences in the reward section

The difference in the intensity of neuronal firing activity between left and right rewards was assessed by comparing the mean firing rate of neuronal activity elicited in the reward section of left and right trials (paired t-test on mean firing rates; *P < 0.05*; Figure 6B column 3).

##### Relationship of encoding preferences in the reward section to those in the remainder of the maze

To assess whether the trajectory-specific firing activity in the maze was linked to discrepancies in the response to left and right rewards, we followed the next steps of analysis. First, for every neuron and every maze section, we calculated the mean value of the relative firing rate difference between left and right trials 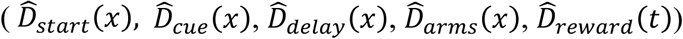. Then, for each neuronal group, we calculated the linear relationship (Pearson’s correlation) between the reward section mean values, to those in the remainder of the maze (Pearson’s correlation; Figure 6C).

#### Immunohistochemistry

After completion of the recording sessions, which lasted about a month, mice were anesthetized with isoflurane and perfused transcardially with 10 ml PBS and 10 ml paraformaldehyde (4%), before they were decapitated. Brains were then removed, post-fixed and coronal slices (100 μm) were prepared. The primary antibodies used were rabbit anti-tyrosine hydroxylase (TH) and chicken anti-GFP. The secondary antibodies used were AlexaFluor 549 anti-rabbit and 488 IgG anti-chicken, respectively. Sections were further stained with DAPI to visualize nuclei. Image acquisition was performed with a fluorescence microscope NanoZoomer (Hamamatsu, Japan) system.

#### Methodological considerations

##### Arranging firing rate by position

With only a handful of exceptions (our report belongs to this minority group), scientific manuscripts reporting the encoding properties of DA neurons arrange neuronal responses by time and align them by key behavioral events, such as trial start, visual cue presentation, reward delivery, etc. We also attempted to arrange firing activities by time, but soon we came to the realization of the inherent caveats of this method in the T-maze task.

The encoding properties of DA neurons have mostly been studied in classical conditioning or reinforcement learning tasks. In those tasks, the animal does not control the exact timing of behavioral events. The experimenter determines when trials begin or end, when visual cues or rewards are delivered, and how much time elapses between cue onset and rewards. Then, usually, post-hoc analysis looks for neuronal responses that are time-aligned to significant task events. For this type of analysis, the researcher should be careful so that the epochs of interest between successive events must not overlap, and most of all, important events are not captured within the epoch of interest of the preceding or succeeding events.

However, in the T-maze task, the timing of behavioral events was completely controlled by the animal, causing a high amount of variability in the timestamps of the task events. For example, within a recording session, animals would consume unique trial rewards within a range of 1 to 10 seconds. Also, the trial duration would usually increase throughout the session as the animals progressively became tired and slower. On top of all, there was high behavioral response variability between animals. Therefore, the increased variability in the timing of events between trials, sessions, and animals caused a significant amount of ambiguity when firing activities were arranged by time and aligned by key task events. Figure S5 shows the average firing activity of individual neurons extracted from a single recording session when neuronal firing was arranged by time (A) and by position (B; the position-arranged firing rate of these particular neurons are presented in Figure 3A, but for convenience to the reader are also shown here). In Figure S5A firing activities for left/right trials were also aligned at the timing of the delay offset (sensor 5). The shadow-colored areas show the range of the timestamps of the other important task events (trial start, cue onset, etc.) across 80-100 trials, relevant to the delay offset. It is evident from this example that deciding the epoch of interest for the memory delay is highly ambiguous.

This important caveat could be easily resolved by arranging firing activities by position. This way, we could produce reliable neuronal firing averages from recording sessions and perform comparisons between behavioral tasks and animals.

##### Identifying and characterizing RPE signals in the Tmaze task

The role of DA in processing RPE signals has been studied extensively with classical conditioning tasks (Ljungberg et al., 1991; Schultz et al., 1993; Schultz et al., 1997; Schultz, 2002; Tobler et al., 2005; Kim et al., 2020). In this Pavlovian paradigm, animals are usually physically restrained and are not trained to make decisions, also, they are exposed to a strictly controlled sensory environment and receive easily accessible and immediate rewards. The present study was designed to investigate the memory-encoding properties of individual neuron in a high-dimensional environment. We did not observe strong manifestations of RPE signaling (Figure S5C). However, compared to classical conditioning studies, in the T-maze task neurons were under the control of numerous behavioral parameters that could be masking cue-related responses, and therefore we cannot draw safe conclusions about the computational role of DA neurons on RPE signals. So far, only manipulating the behavioral parameters with virtual reality tools, has provided insight into the RPE-related responses of DA neurons in goal-directed behavioral tasks like ours (Kim et al., 2020).

## Acknowledgments

This study was supported by the Ministry of Education, Culture, Sports, Science, and Technology (Grants-in-Aid for Scientific Research 18H02711 and 18H05525), the Mitsubishi Foundation, the Naito Foundation, and the Japan Agency for Medical Research and Development (AMED).

In memory of Miles Adrian Whittington. A true mentor and friend.

## Author contributions

V.G. and S.F. designed the experiments. V.G. performed the experiments and analyzed the data. V.G. and S.F. wrote the paper.

## Supplementary Materials for

**This PDF file includes:**

- Supplementary Text 1
- Figs 1–11
- Table 1

### Supplementary Texts

**Supplementary Text 1. Evaluating the contribution of behavioral variables in trajectory-specific activity with multiple regression analysis**

We designed a generalized linear regression model in which the dependent variable was the position-arranged firing rate, and the independent variables (predictors) were the animal’s running speed, trial number, performance rate (cumulative correct rate), current trial accuracy (reward or not), previous trial accuracy (reward history), and lap trajectory (left or right). Since the permutation analysis of the original spike trains revealed that the trajectory-specific effect on neuronal firing activity was highly dependent on the animal’s position (occurring mainly in the delay and side-arm sections), we included the joint effect of lap trajectory and position in the training model instead of testing the effect of trajectory alone. A major advantage of regression analysis is that one can dissociate the inherently bound effects of the independent variables. For example, we can examine the influence of the lap trajectory variable on neuronal firing activity without including the effect of the running speed variable (which could potentially differ between left and right trials).

For each neuron, we produced model predictions for the correct left and right trial average firing rates (Figure. S7A, dashed lines) from which we calculated the predicted firing rate difference. We subsequently examined the contribution of each independent variable to the trajectory-specific firing activity by shuffling the trajectory labels assigned only to this particular variable. If the tested variable exerts a significant effect on the firing rate, then shuffling the trajectory labels would produce a significant reduction in the predicted firing rate difference. We examined the same pool of neurons that we reported on the memory task, as shown in Figure 3B (n=104 DA and n=74 GABA-identified neurons). In the delay region, the joint effect of lap trajectory and position (trajectory × position predictor) contributed significantly to the predicted average rate difference of 23 DA neurons (Figure S7B and S7C top), 17 of which overlapped with the 22 trajectory-specific neurons identified with the permutation analysis (Figure S7C bottom). Of the GABA neurons, 34 were modulated by trajectory and position (Figure S7B and S7C top), 30 of which were also trajectory-specific (Figure S7C bottom). These results confirm that the trajectory factor was responsible for the firing rate difference shown in Figure 3B. The speed variable significantly modulated 24 DA and 22 GABA neurons (Figure S7B and S7C top), but only four DA and four GABA neurons were co-modulated by trajectory (Figure S7C bottom). The performance and accuracy variables modulated smaller numbers of neurons (Figure S7B and S7C top), and the reward outcome of the previous trial did not co-modulate any of the trajectory-specific DA and GABA neurons (Figure S7C bottom). The trial variable modulated 30 DA neurons and 32 GABA neurons, co-modulating with the trajectory variable of 8 DA and 15 GABA neurons. However, the distribution of the trial predictor coefficient did not differ from a distribution with a mean equal to zero (one-sample *t*-test, *P* > 0.05, Figure S7D), indicating that the effect of successive trials on firing rate did not reflect cognitive processing, but was caused by mechanical reasons; due to the animal’s movements, the distance of the recording channel from the targeted neurons changed continuously, which affected the signal-to-noise ratio and eventually spike detection. In agreement with Engelhard et al., a notable proportion of DA neurons (36%) and GABA neurons (74%) were co-modulated by more than one behavioral variable Figure S7E (Engelhard et al., 2019).

Overall, the regression analysis confirmed the results of the permutation analysis regarding the significant effect of trajectory on midbrain neuronal activity during memory-dependent decisions. In addition, it demonstrated that the remaining independent variables included in our model cannot fully explain trajectory-specific firing activities in the delay period of the memory task.

### Supplementary Figures

**Figure S1.**
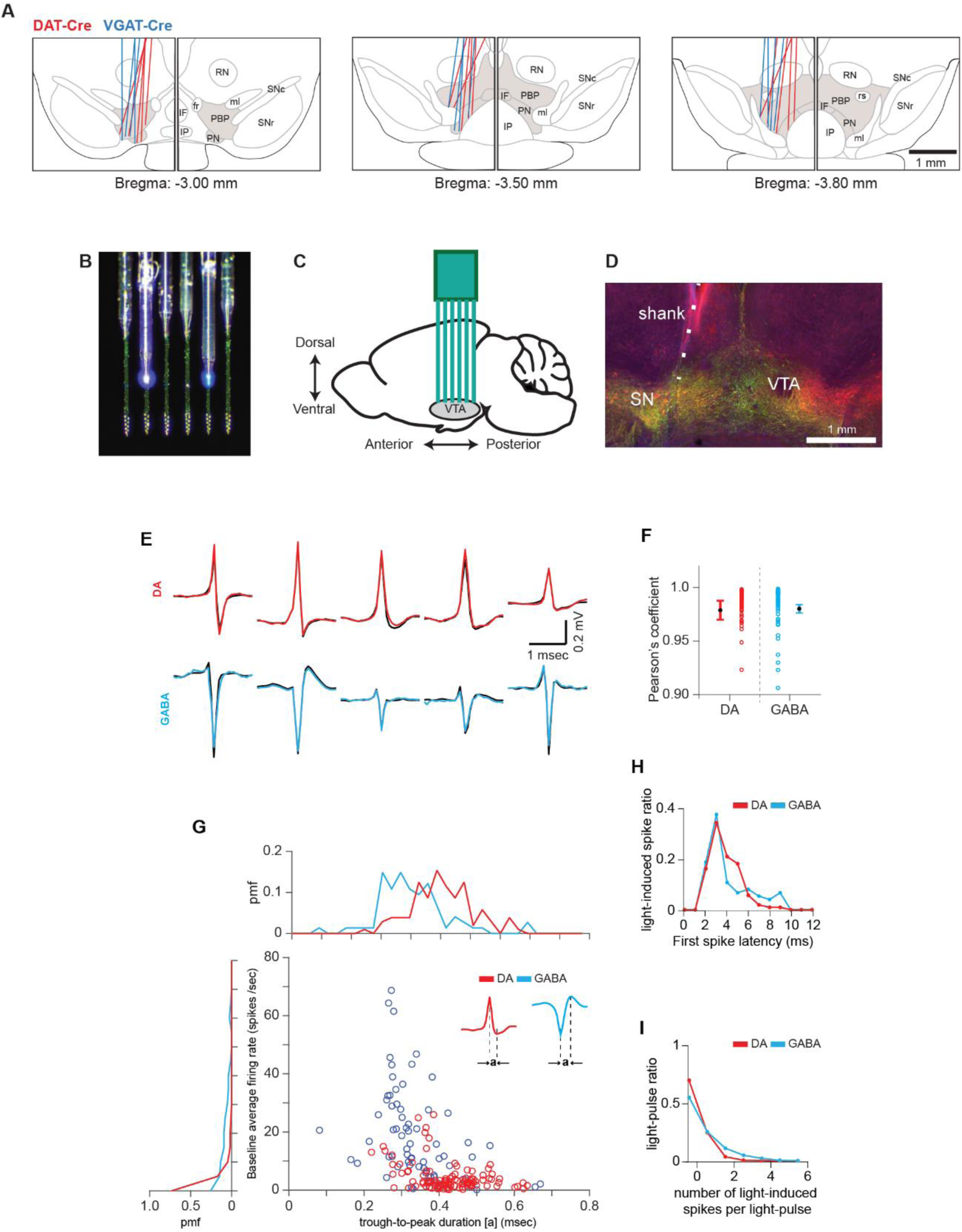
Validation of optogenetic identification results. Related to Figure 1. **(A)** Serial reconstructions of recording sites along the rostro-caudal axis of coronal midbrain slices for DAT-Cre (red) and VGAT-Cre (blue) animals. *fr*: fasciculus retroflexus, IF: interfascicular nucleus, IP: interpeduncular nucleus, *ml*: medial lemniscus, PBP: parabrachial pigmented nucleus, PN: paranigral nucleus, RN: red nucleus, *rs*: rubrospinal tract, SNc: substantia nigra pars compacta, SNr: substantia nigra pars reticulata. **(B)** Photograph of the silicon probe with attached optical fibers. Optical fibers (core diameter 50 μm) were glued to the shanks to ensure a firm, accurate, and durable insertion into the deep midbrain. Optical fibers on shanks 2 and 5 were coupled to blue laser diodes (450 nm), maintaining a certain distance from the tip of the shank to ensure that blue light could irradiate the whole span of the recording electrode array. **(C)** Schematic drawing of the mouse brain in the sagittal plane, illustrating the insertion of the silicon probe into the VTA. **(D)** Immunofluorescence microscopy image of a coronal brain slice of a DAT-Cre mouse injected with the optogenetic virus in the left VTA. DAT: dopamine transporter (red); YFP: yellow fluorescent protein (green); SN: substantia nigra. NOTE: the white dashed line highlights the trace by the probe shank on the tissue. **(E)** Mean waveforms of light-induced (colored) and spontaneous (black) spikes discharged by identified DA (red) and GABA (blue) neurons in a single light stimulation session. **(F)** Pearson’s correlation coefficient between the waveforms of spontaneous and light-induced spikes for the identified DA (red) and GABA (blue) neurons. The light-induced spike waveforms were almost identical to the spontaneous waveforms for all the identified neurons. This was reflected by the high mean correlation coefficient (mean ± standard error of the mean; DA: r = 0.98 ± 0.09, *P < 0.05*; GABA: r = 0.98 ± 0.03, *P < 0.05*). **(G)** Trough-to-peak spike duration vs. average spontaneous firing rate raster plot for the identified DA (red) and GABA (blue) neurons. The side plots show the distributions of the X- and Y-axis values in the main plot (pmf: probability mass function). In agreement with earlier reports (2) DA neurons fired spikes with lower rates (DA: 4.50 ± 5.00 spikes/sec, GABA: 17.34 ± 15.72 spikes/sec; unpaired t-test comparing the spontaneous rates: t_(176)_ = 7.80, *P* < 0.001) and wider waveforms (DA: 0.46 ± 0.08 ms, GABA: 0.34 ± 0.10 ms; unpaired t-test comparing trough-to-peak values: t_(176)_ = 5.96, *P* < 0.001) than the GABA neurons. **(H)** Latency of the first spike discharged during light stimulation for the identified DA (red) and GABA (blue) neurons. **(I)** Number of light-induced spikes per light pulse for DA (red) and GABA (blue) neurons.

**Figure S2.**
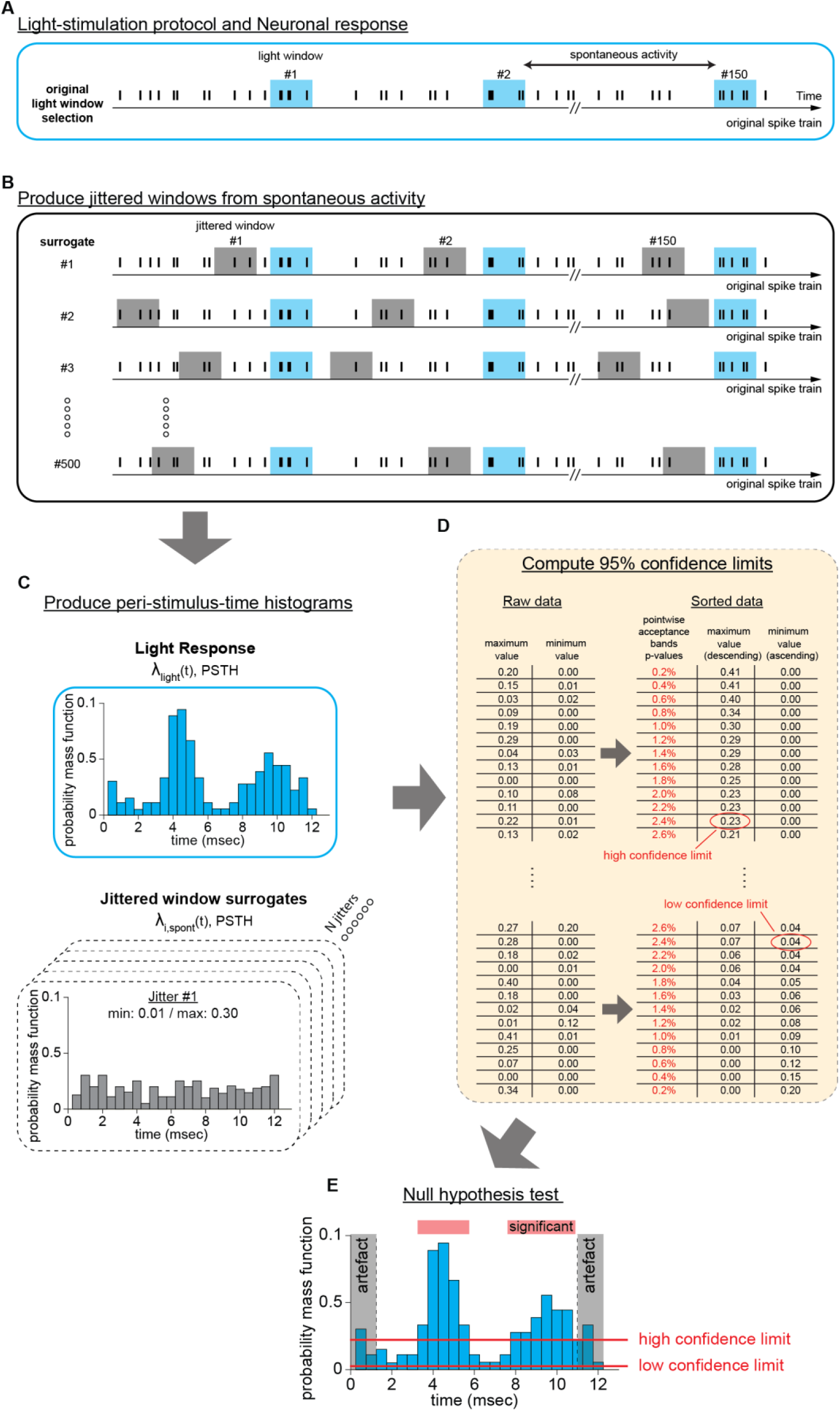
Statistical method for optogenetic identification. Related to Figure 1. To identify the light-responsive units, we compared the light-induced average responses to average spontaneous firing activity. To this end, we used the following jittering method. **(A)** Schematic representation of a spike train during a light stimulation session. Vertical black bars represent spiking events. The blue shaded areas indicate the light-stimulation windows. The firing activity between light stimulations is spontaneous. **(B)** To identify light-responsive units we raised the hypothesis that the relative firing rate during light stimulation, ^*λ̂*^_*lig*ℎ*t*_(*t*), did not differ from the spontaneous firing rate, ^*λ̂*^_*spont*_(*t*). To do so, we employed a jittering method in which a randomly jittered window (12 ms) fitted in the spontaneous activity period preceding the light-pulse (that is, the jittered window onset never preceded the offset of the preceding light-pulse and the offset never succeeded the onset of the succeeding light pulse). We produced as many jittered windows as the light-pulses and we repeated this process 500 times in total. **(C)** For every neuron we produced peri-stimulus-time-histograms (PSTHs, 12 ms, 1 ms bins) of the relative firing rate, by superimposing the spike trains of every light stimulation pulse. If *n*^(*k*)^(*t*) is the number of spikes of a single neuron in post-light onset time *t* of the *k_th_* stimulation pulse, then, the average firing rate, 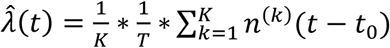, where *K* is the number of stimulation pulses, *T* is the bin size (1 ms) and *t*_0_ the light-pulse onset time, represents the relative firing rate at time *t*. In the end we produced one ^*λ̂*^_*lig*ℎ*t*_(*t*) and 500 ^*λ̂*^_*spont*_(*t*) firing rates. **(D)** From every, ^*λ̂*^_*i*,*spont*_(*t*) histogram we extracted the maximum and minimum values and stored them in two separate vectors. Vectors were sorted, and from them, the 95% confidence limits were calculated. These were the 2.5% largest value from the maximum PSTH vector and the 2.5% smallest value from the minimum PSTH vector (that is 13rd from the 500 rows). Units with ^*λ̂*^_*lig*ℎ*t*_(*t*) PSTH exceeding at any time point (between 1 ms and 11ms) the high confidence limit, were identified as light-responsive. **(E)** If the ^*λ̂*^_*lig*ℎ*t*_(*t*) PSTH fell below the low confidence limit at any time point, then the unit was characterized as light-inhibited due to synaptic transmission (we did not find any light-inhibited units). Otherwise, those units whose PSTH did not violate any of the confidence limits were characterized as unidentified.

**Figure S3.**
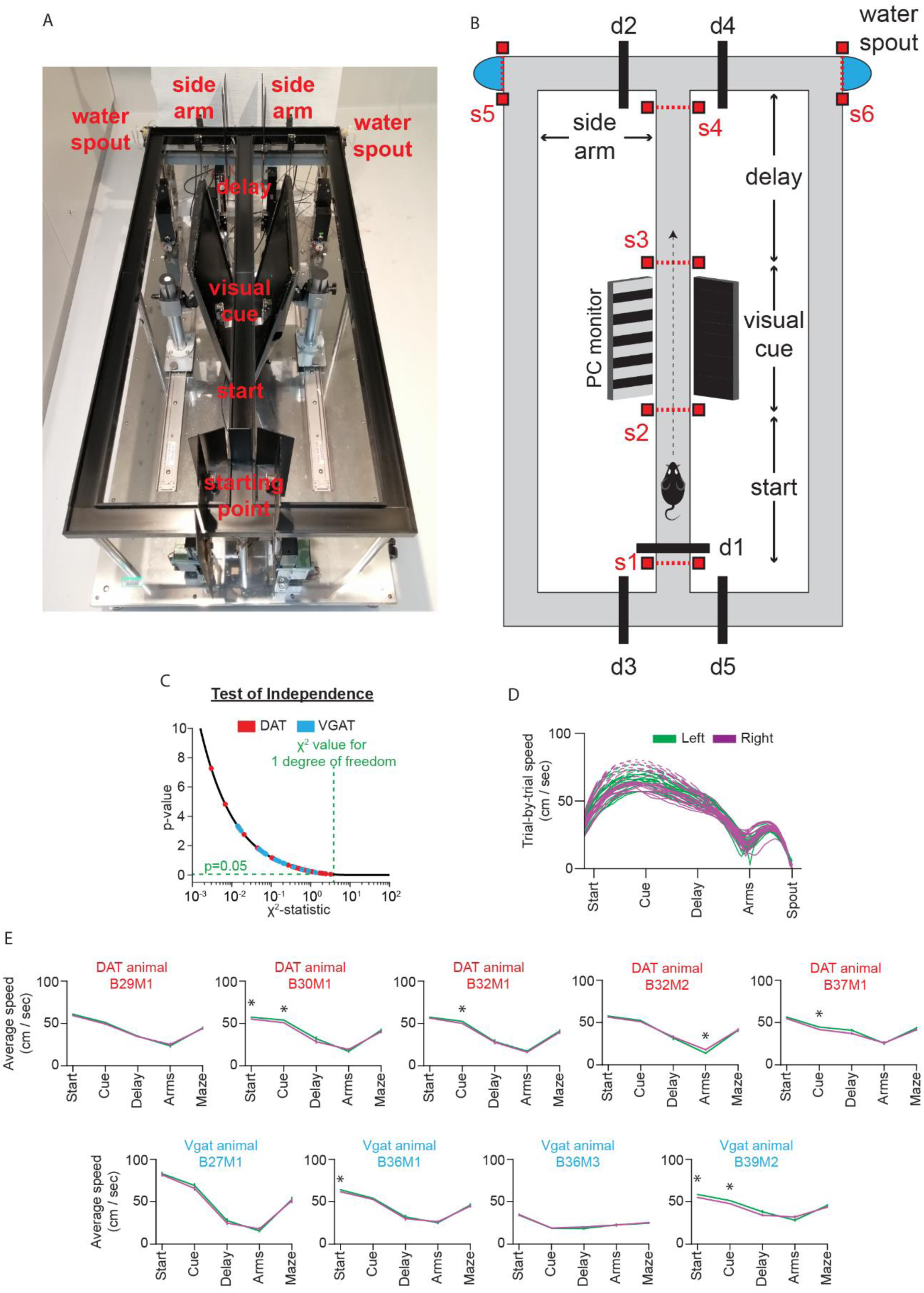
T-maze configuration, behavior and running speed assessment in the memory task. Related to Figures 2, 3, 4, and 5. **(A)** Photograph of the T-maze apparatus with maze sections labeled. The mice were trained on different variations of an associative T-maze task with visual instruction cues. The behavioural apparatus was a Figure-eight T-maze (O’Hara & CO., LTD, Tokyo, Japan), 50 cm tall, with a main arm (120 cm) and side arms (30 cm each). Infrared light-beam sensors placed at key position points on the maze defined the beginning and end of the successive maze sections. The apparatus was equipped with sliding doors, which prevented the animals from developing unwanted behavior (e.g. moving backwards). Sensor activations were on-line monitored, and significant behavioural events (e.g. visual-cue presentation, reward delivery, door opening/closing) were dictated by an in-house behavioural software written in MATLAB (Mathworks, MA, USA), through a multi-signal processor interface system (RX6; Tucker-Davis Technologies, FL, USA). The moment-by-moment position of the animal on the maze apparatus was monitored continuously by a video tracking system, composed of a red light-emitting diode mounted on the mouse head stage and a video camera (39 frames/sec) hung by the room ceiling. Video data were stored for offline analysis. **(B)** Detailed schematic representation of the T-maze apparatus. Infrared light-beam sensors (s1–s6) were mounted securely at key positions on the maze to define the beginning and end of the maze sections (start: s1–s2; visual cue: s2–s3; delay: s3–s4; side-arms: s4–s5/s6; reward: s5/s6). The first activation of s5 or s6 sensors (i.e., 1^st^ lick) triggered the water-delivery pump. Five sliding doors restrained the animals from unwanted moves. Every trial began with the animal activating s1. At this point, the doors d1, d3, and d5 remained closed, isolating the animal in the starting location for approximately 2 s. This served two purposes: (1) to drain the waterspouts and (2) to prevent the animal from making decisions on the next trial guided by the previous trial outcome. After door d1 opened, the animal could run freely along the main arm. Activating s2 and s3 resulted in the visual cue onset and offset, respectively. Activating s5 triggered the left water-delivery pump, and door d3 opened while d1 closed. Activating s6 triggered the right water-delivery pump, and door d5 opened while d1 closed. In the memory task, doors d2 and d4 were always open. In the no-cue-no-choice task, doors d2 and d4 were initially closed. However, when the mouse activated s4, one door opened pseudo-randomly. **(C)** To assess the influence of behavioural left-right biasing in decision making we applied the chi square (χ^2^) test of independence. The null hypothesis was that correct performance was independent of behavioural choices. Black line illustrates the chi-squared distribution for 1 degree of freedom, along with the χ^2^ values from the independence test for every session (DAT-Cre: red dots and VGAT-Cre: blue dots). The test was passed in every session (the level of significance was set to 5%, corresponding to χ^2^ = 3.84 for 1 degree of freedom). **(D)** Representative example of trial speeds (cm/sec) in a single recording session of the memory task. **(E)** Shown are running speeds (mean ± standard error of the mean) categorized by maze sections and averaged across sessions for left (green) and right (magenta) trials for individual VGAT-Cre and DAT-Cre mice. Although differences were small, they occasionally reached significance. Importantly, we did not detect speed differences between left and right trials in the delay section (unpaired t-test between left and right speed per region, * *P* < 0.05).

**Figure S4.**
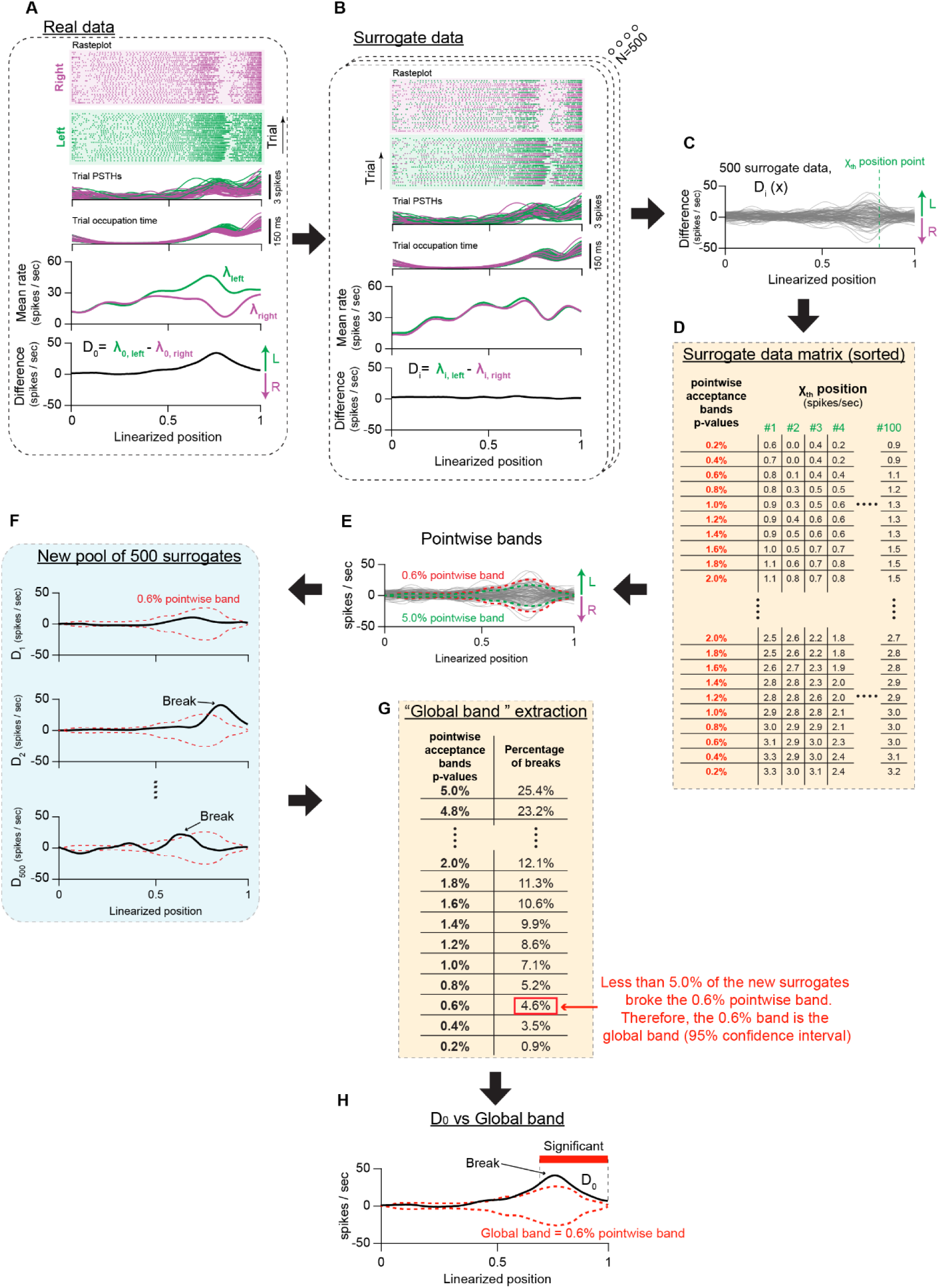
Statistical method for identifying trajectory-specific neurons (permutation method). Related to Figure 3. We applied a permutation method reported elsewhere (Fujisawa et al., 2008) to identify neurons with trajectory-specific encoding properties. The motivation behind this analysis is that if the lap trajectory contributes to the firing rate difference observed at certain positions, then shuffling the trajectory labels assigned to the spike trains of individual trials would cause a marked reduction in the rate difference. This process is described below. **(A)** From the spike trains of the correct left and right trials (top; raster plot), we estimated the trial relative firing rates (^*λ̂*^_*Left*_(*x*) and ^*λ̂*^_*Rig*ℎ*t*_(*x*); middle). Importantly, we considered the variability in occupation time between positions and trials. Hence, to calculate the firing rates, we divided the number of spikes occurring in every position and trial (trial PSTHs) by the occupation time in the same position and for the same trial (trial occupation time). We subsequently produced the original average rate difference (*D*_0_(*x*), bottom) between the left- and right-correct trials. **(B)** We shuffled the trajectory labels assigned to the trial spike trains along with the respective occupation times and re-calculated the difference *D*_*i*_(*x*) of the permuted labels. **(C)** We produced 500 surrogates with permuted differences *D*_*i*_(*x*). In the end, we created a 500-by-100 matrix (shuffles-by-position) of *D*_*i*_(*x*) surrogates. **(D, E)** From these surrogates we extracted the 95% confidence interval (lower and upper confidence limits) for the null hypothesis test. To do so, first, we calculated the confidence interval of each position point *x*. The difference values *D*_*i*_(*x*) in every position (column) were sorted and every row in the new matrix was treated as a potential confidence limit with different *P*-value. We term it as “pointwise acceptance band”. If the original data *D*_0_(*x*) breaks the pointwise band, it corresponds to rejecting the null hypothesis at position *x*. However, if this procedure is repeated for every position *x* (i.e., 100 position points) in the maze, it raises the issue of multiple comparisons. **(F, G)** To address this, we constructed the “global band”, which can control errors of any false rejection across multiple indices. We produced another 500 permutations and calculated the percentage of the surrogate data *D*_*j*_(*x*) that broke the pointwise band candidates at any position points. If the percentage was more than 5%, we replaced the pointwise band with lower P-value. We repeated this process until a pointwise band candidate was exceeded by less than 5% of the new surrogates. When this happened, the pointwise band was used as the “global band” (i.e., 95% confidence interval) for the hypothesis test. **(H)** The null hypothesis was rejected if the original difference *D*_0_(*x*), exceeded at any position *x* the global band. Then, the spatial extent of significance was defined by the number of position points exceeding the 95% confidence interval.

**Figure S5.**
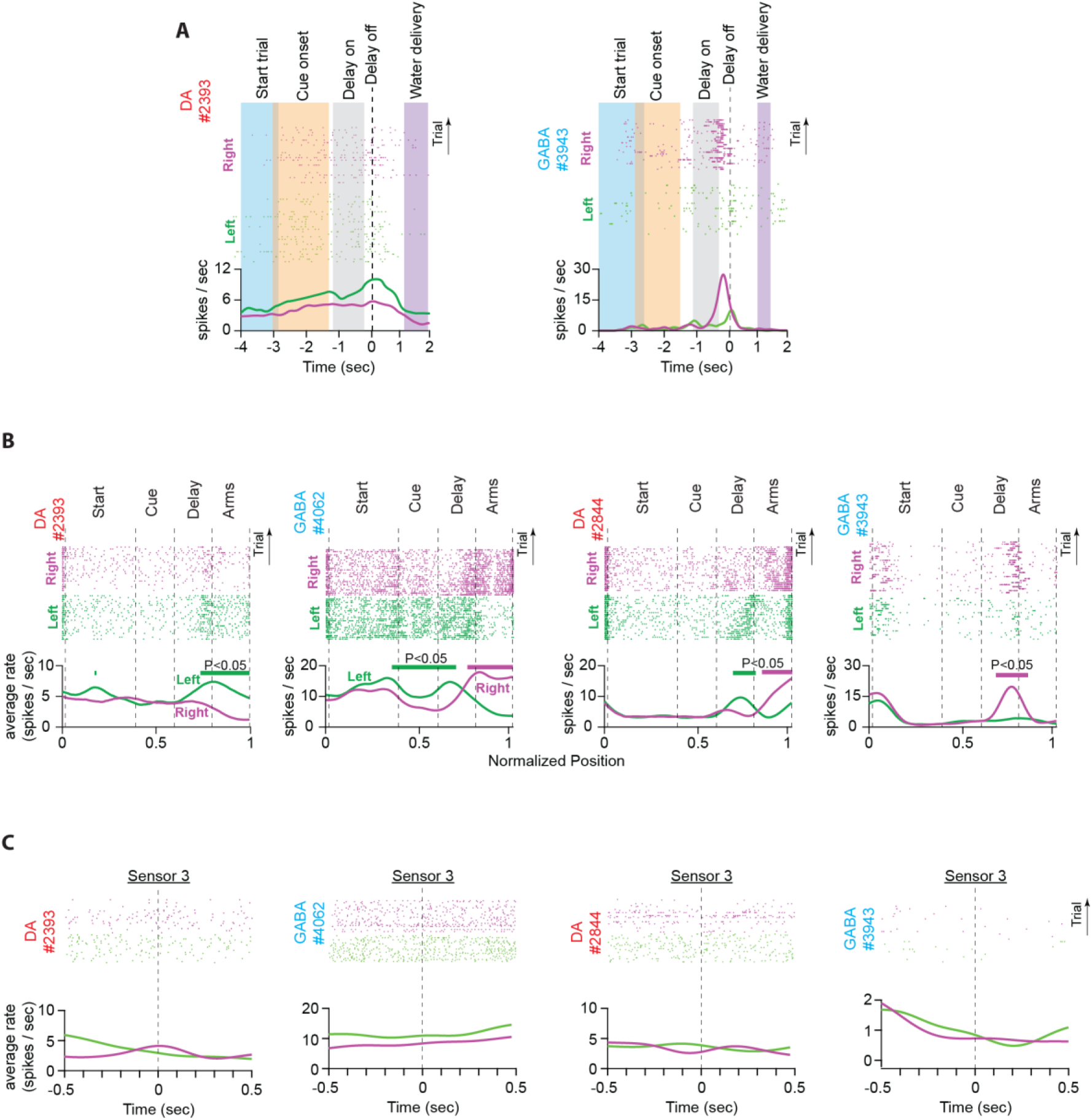
Arranging neuronal firing activities by time or position. **(A)** Firing activities of representative neurons (DA#2393 and GABA#3943) arranged by time and aligned at the offset of memory delay. The colored bands illustrate the range of the timestamps of key task events across the trials of a single recording session. Timestamps differed significantly across trials, sessions, behavioral protocols, and animals. As a result, we could not define a fixed epoch for every maze section with adequate duration to analyze the neuronal responses. **(B)** Firing activities of representative neurons (including DA#2393 and GABA#3943) arranged by position (same plots as in figure 3A). Notably, the position of key task events was the same across trials, sessions, protocols and animals, enabling the comparison of neuronal responses between animals and behavioral protocols. **(C)** Firing activities of the same neurons as in Figure S5B, arranged by time and aligned by the timestamps of sensor 3 crossings. Consistent with earlier reports (Howe et al., 2013; Kim et al., 2020), while animals navigated the maze, receiving a continuous sensory input and making accurate estimations of the timing of key task events, we did not observe profound discharge rate elevations in response to the visual cue onset that would resemble strong RPE signaling.

**Figure S6.**
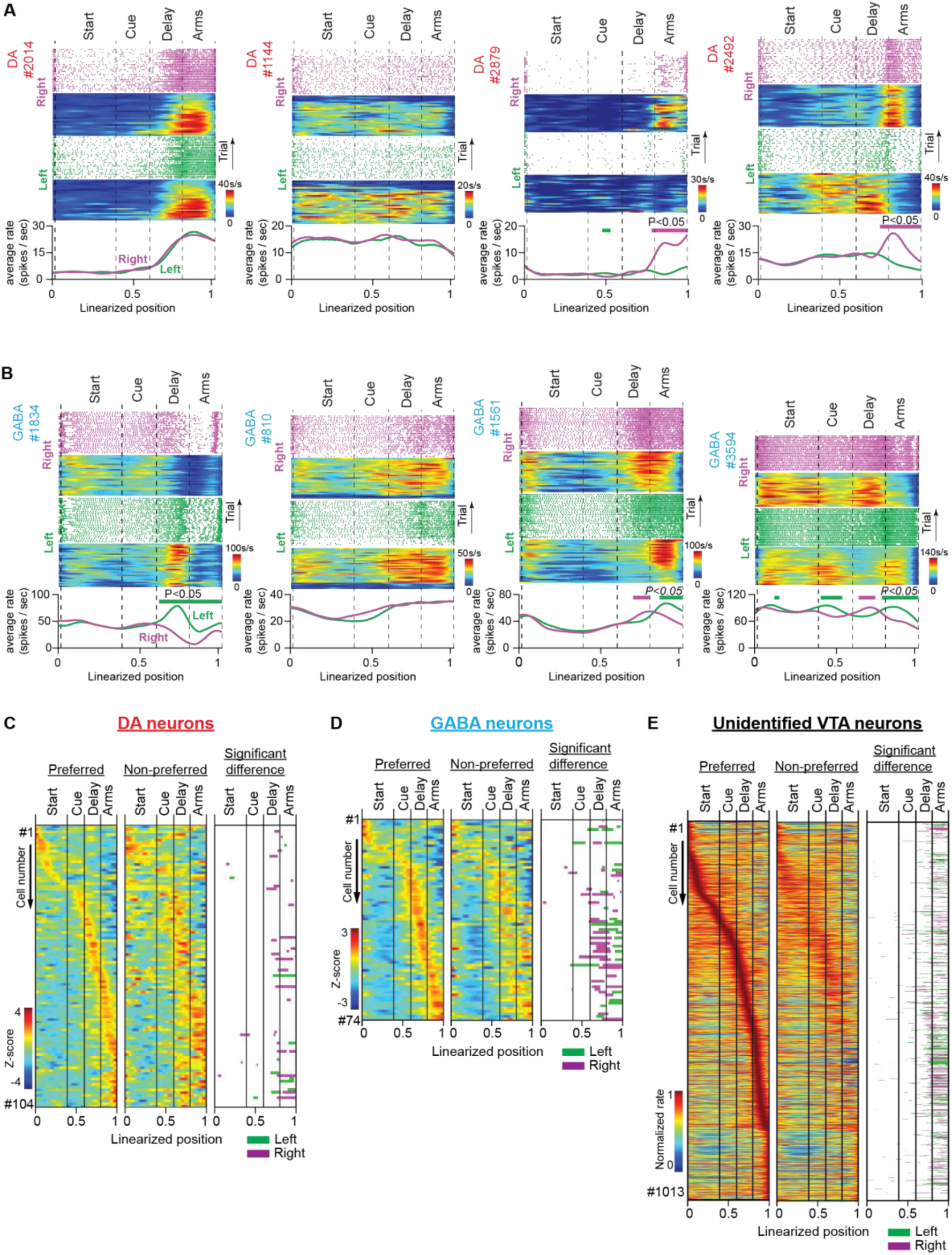
Firing activities of midbrain DA and GABA neurons in the memory task. Related to Figure 3. All plots included in this figure were obtained from data recorded in the memory task. **(A and B)** Representative discharge activity of DA (a) and GABA (b) neurons. In each example: (Top) Raster plots of the spikes, arranged by trial, and their corresponding firing rate heatmaps as a function of position in right (purple) and left (green) trials. (Bottom) Average firing rates for correct left and right trials. Note that the firing rate (spikes/s) is plotted as a function of position but has been normalized by the amount of time the mouse occupied each position in every trial. The thick lines above the firing rates represent segments with significantly different firing rates between the correct right and left trials. **(C and D)** Heatmaps of the standardized average firing rates for the preferred (first column) and non-preferred (second column) trajectories of DA (c) and GABA (d) neurons. These plots show the number of standard deviations by which the average firing rate varied above or below the mean rate of correct trials as a function of the position on the maze. The row arrangement was the same as that in Figure 3B. The third column of the significant firing-rate difference is the same as that in the third column in Figure 3B. **(E)** Heatmaps of the normalized average firing rate for preferred (first column) and non-preferred (second column) trajectories and the significant difference between them (third column) for the unidentified VTA neurons.

**Figure S7.**
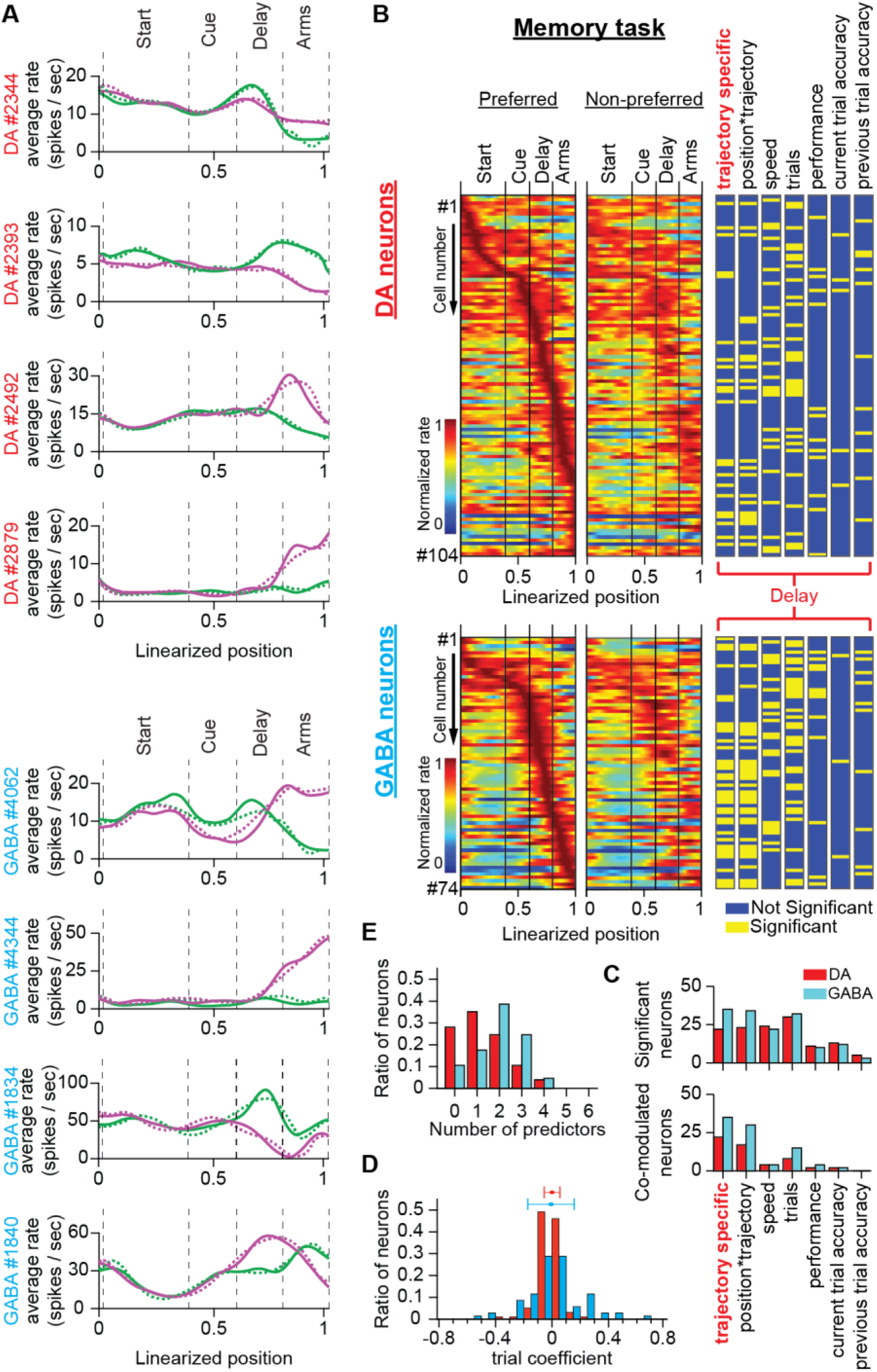
Evaluating DA and GABA neuronal responses to specific behavioral variables in the memory task through regression analysis. Related to Figure 3. **(A)** Average firing rates of representative DA and GABA neurons extracted from real firing rate data (solid lines) and encoding model predictions (dashed lines) for left (green) and right (magenta) trials. **(B)** (Left and Middle columns) Normalized firing rate heatmaps for preferred and non-preferred trajectories for DA and GABA neurons recorded in the memory task (the same as the ones in Figure 3B). (Right column) Significant contributions in the firing rate difference (yellow lines) by individual behavioral variables in the memory delay extracted with the permutation analysis of the encoding model predictions. **(C)** (Top) Number of neurons significantly modulated by one of the independent variables. (Bottom) Number of neurons co-modulated by trajectory (trajectory-specific) and one of the individual variables. (Note: In B and C, the trajectory-specific neurons correspond to the significant neurons extracted using the permutation analysis of the original firing rates; third column in Figure 3B). **(D)** The distribution of the trial coefficient for DA and GABA neurons along with the mean ± standard deviation values. **(E)** Histogram of the number of independent behavioral variables co-modulating neurons in the delay region.

**Figure S8.**
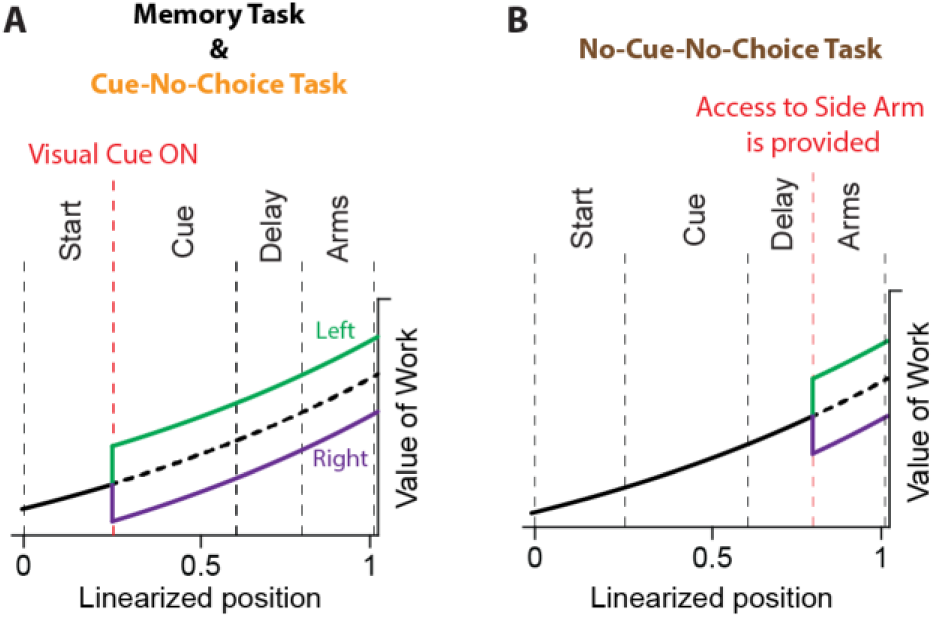
The dopamine signaling model of incentive motivational drive. Related to Figures 2, 3 and 4. Striatal DA levels ramp up when mice navigate a maze or corridor in search for reward (Howe et al., 2013; Hamid et al., 2016; Kim et al., 2020). Initially, this evidence challenged theoretical neuroscientists, since the time-course of this ramping activity (lasting a few seconds) did not conform either with the prediction-error (RPE, phasic activity) or reward rate (tonic activity) theories (Niv, 2013). However, further research on this topic, provided empirical evidence to support a theory that DA conveys a signal for incentive motivational drive in the form of state-action value (value of work; (Hamid et al., 2016), or its derivative RPE; (Kim et al., 2020)). Accordingly, the motivational value of future rewards is exponentially discounted with time or distance (the theory was developed on an adaptive decision-making framework; Hence the value of reward is defined by reward probability; figure 4 in (Hamid et al., 2016)). Different reward values correspond to different functions of discounted state-action values. If a cue predicts a reward with higher value, then the state-action value jumps to the discounted value function of that reward. In our study, key parameters that could potentially influence incentive motivational drives were equal between left and right trials (reward amount, lap trajectory distance, etc.) and moreover, behavioral performance did not indicate choice biasing (Figures S3C). However, we cannot rule out the possibility that left and right rewards were valued differently, producing distinct value functions. If this is true, then the trajectory-specific activities elicited by subgroups of DA and GABA neurons could reflect differences in the state-action values for left and right trials, and not memory-dependent decisions. **(A)** The plot illustrates the hypothesis of different discounted value functions (i.e., the state-action value is a function of position and trajectory). In the memory task and the cue-no-choice task the assignment of the state-action value function takes place when the visual cue signals which reward is available. **(B)** However, in the no-cue-no-choice task, the assignment occurs only when the mice are given access to one of the side arms.

**Figure S9.**
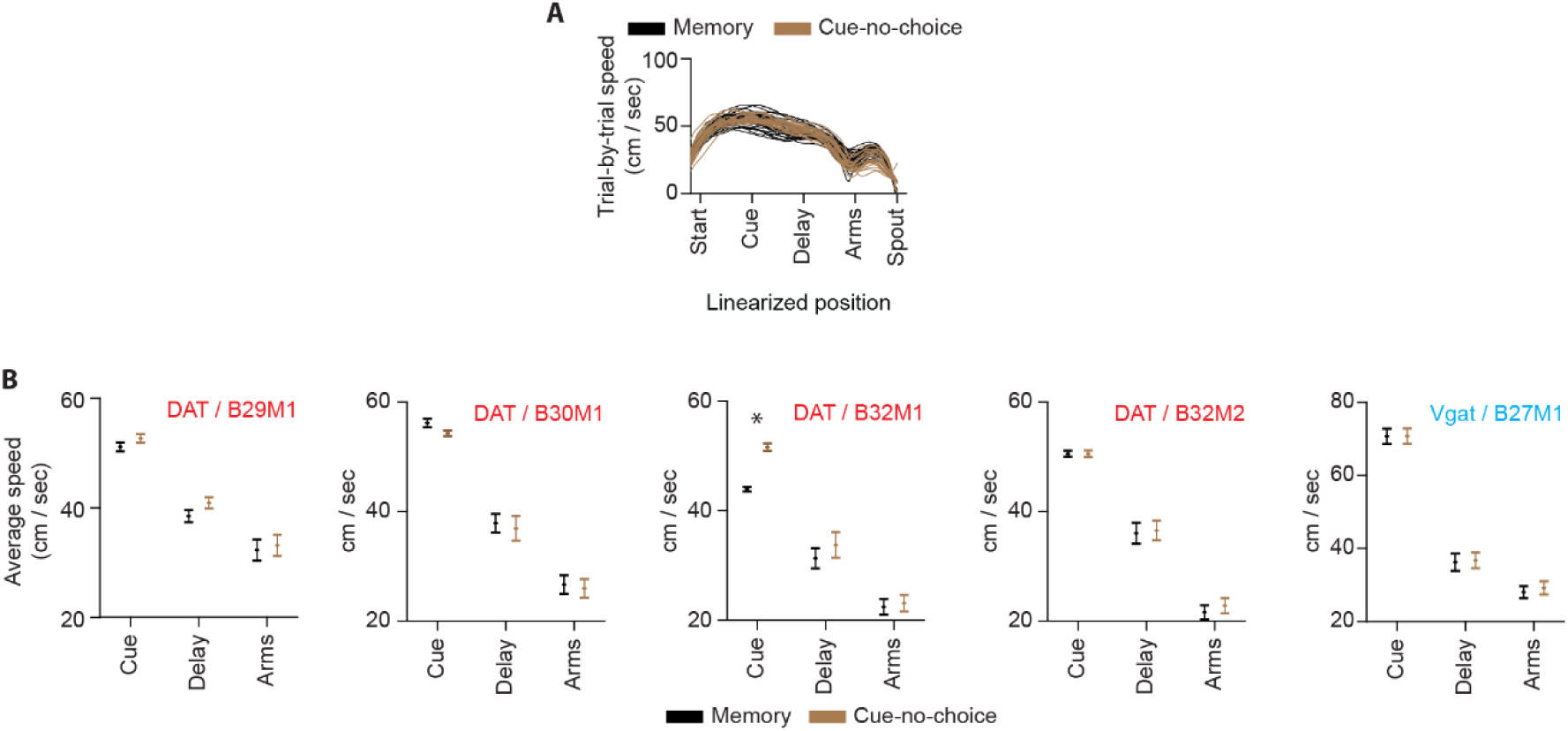
Running speed differences between the memory task and the cue-no-choice task. Related to Figures 3 and 5. **(A)** Representative example of running speeds (cm/s) for memory (black) and cue-no-choice (orange) trials in a single session. Speed differences were evident at the T-intersection (between delay and arms). **(B)** Average running speed values (mean ± standard error of the mean) grouped by maze section and behavioral task for each of the DAT-Cre and VGAT-Cre animals performing the memory and cue-no-choice tasks. Importantly, no differences were detected in the delay and side-arm sections between memory (black) and cue-no-choice (brown) task trials (* *P <* 0.05, unpaired t-test on speed).

**Figure S10.**
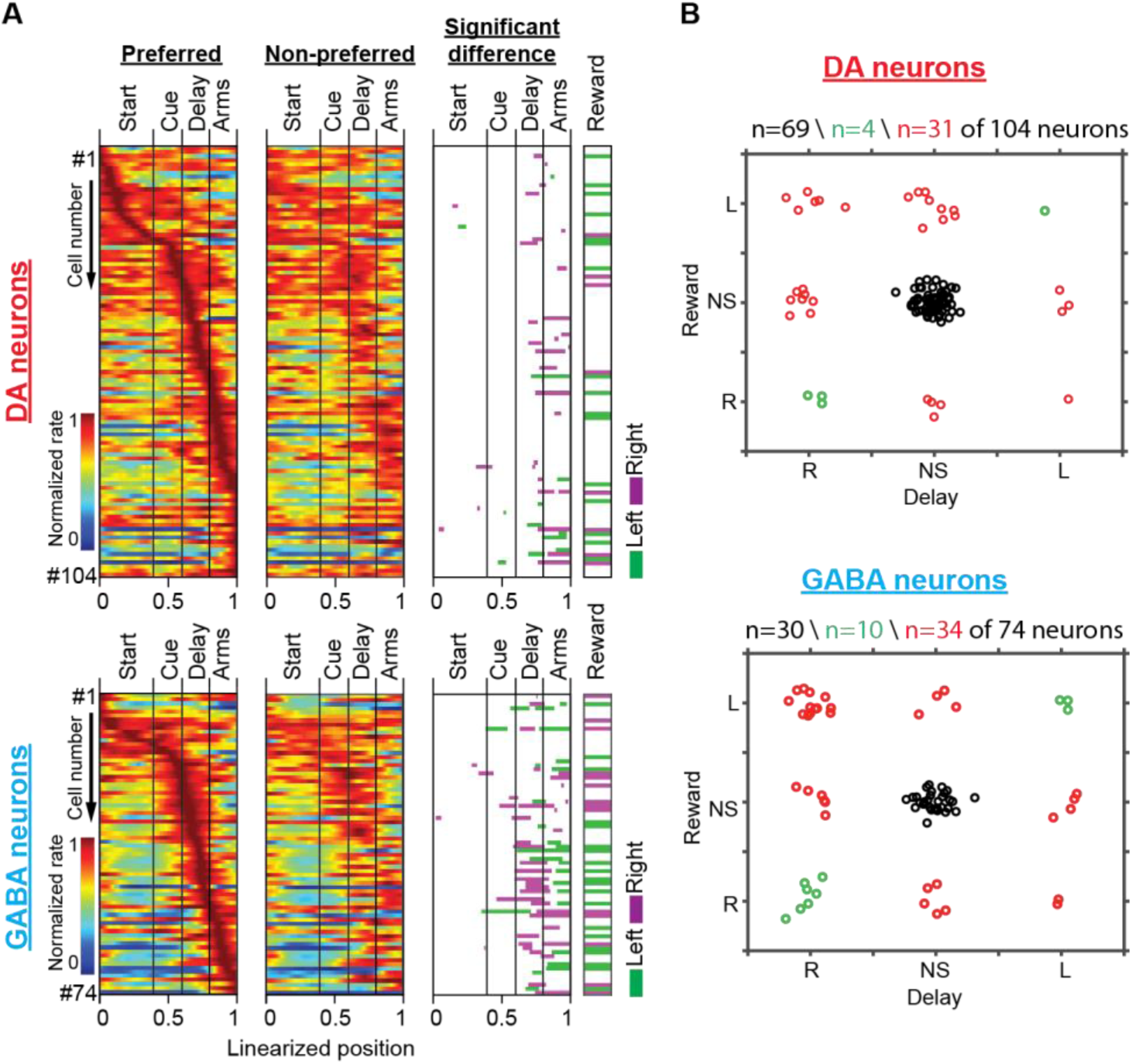
DA neurons which encode memory information for a specific trajectory do not show preference for the same trajectory reward. **(A)** Heatmaps of neuronal population responses organized by preferred lap trajectory (first column) and non-preferred lap trajectory (second column) for DA neurons (Top; *n* = 104 units, 35 sessions in five mice) and GABA neurons (Bottom; *n* = 74 units, 25 sessions in four mice). The third column shows maze segments with significantly different discharge rates between preferred and non-preferred trajectories for the start, visual cue, delay, and side arms sections. The fourth column shows neurons with significant discrepancies between the left and right reward-related responses (paired *t*-test for mean firing rates, *P* < 0.05). Note: The first three heatmaps are adopted from Figure 3B (position-aligned firing rate) and the fourth heatmap is adopted from Figure 6B (third column; time-aligned firing rate). In all the heatmaps, each row contains responses of the same neuron. **(B)** Scatterplots created by dividing neuronal responses from the third and fourth column heatmaps in plot **(A)** into six (6) categories, grouped by the trajectory-specific preference in the delay section (x-axis) and during reward consumption (y-axis). “L” corresponds to a significant preference for the left trajectory, “R” for the right trajectory, and “NS” for non-significant firing rate difference (i.e. no preference). **Note:** to yield a better visual sense of how many observations belong to every category, we randomly jittered each point along the x- and y-axis. In both neuronal populations, more neurons showed opposite significant lap-trajectory preferences (red clusters) between delay and reward sections, as opposed to a small minority of neurons that elicited the same lap-trajectory preference (green clusters). Notably, in agreement with earlier studies reporting preferential contralateral responses of DA neurons (Kim et al., 2015; Parker et al., 2016; Engelhard et al., 2019; Lee et al., 2019; Moss et al., 2020), the majority of memory-specific neurons exhibited a preference for the contralateral lap trajectory to that of the recording site (left hemisphere). Accordingly, seventeen (17) of the twenty-three (23) DA neurons with trajectory-specific activities in the delay period of the memory task, elicited a significant preference for the right trials and only five (5) of them for the left trials. In GABA neurons, the percentage was similar with twenty-six (26) neurons showing a clear preference to the right trials and nine (9) to the left trials.

**Figure S11.**
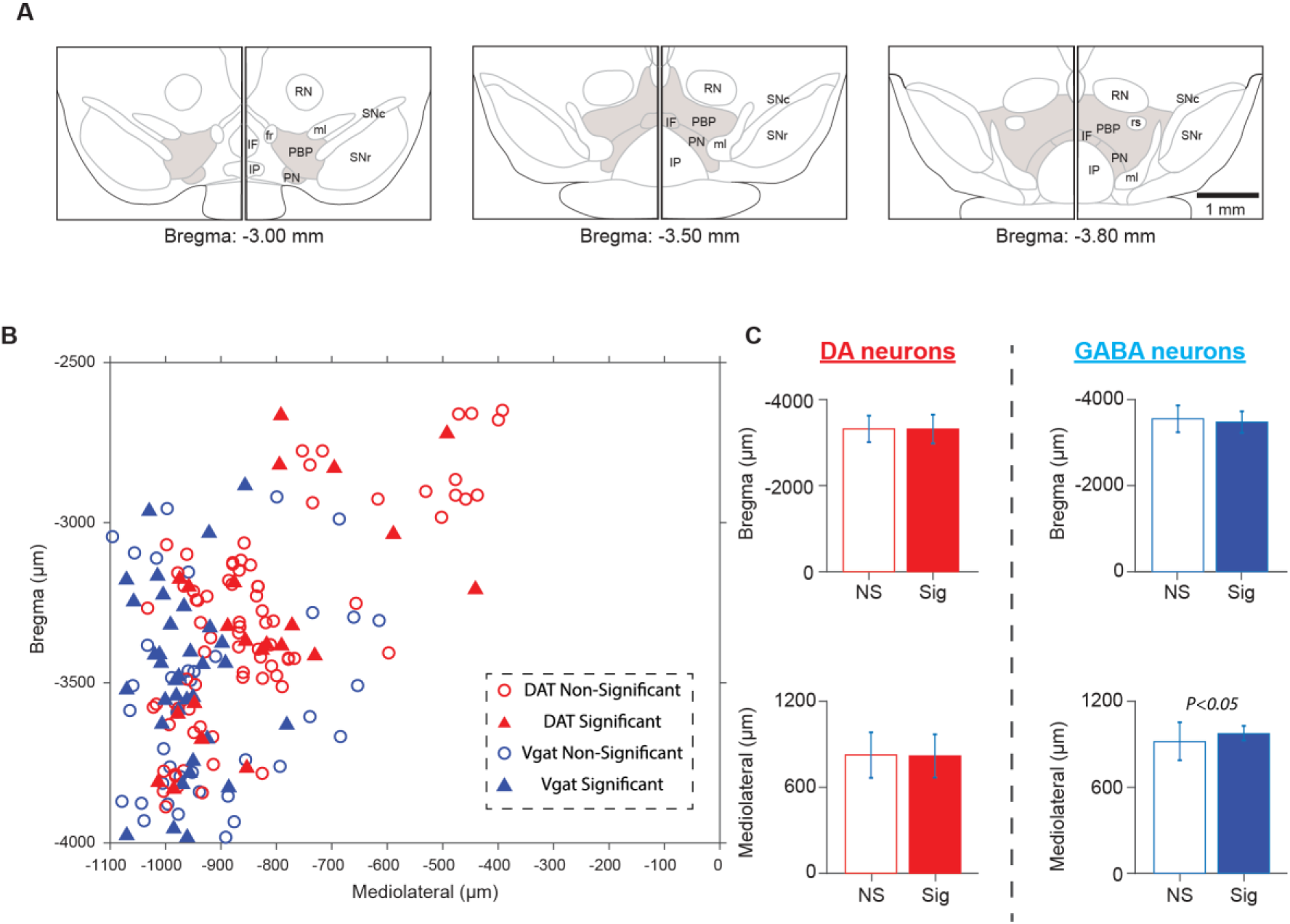
Anatomical organization of memory-specific VTA neurons. (A) Schematic representations of coronal slices containing the ventral tegmental area at different Bregma coordinates. Representations were adopted and modified from The Mouse Brain Atlas in Stereotaxic Coordinates (ref). (B) The approximate stereotaxic coordinates of the optogenetically identified DA and GABA neurons were extracted from the estimated location of the recording channels. From those, a scatterplot was produced. Solid triangles correspond to DA (red; DAT-significant) and GABA (blue; Vgat-significant) neurons with trajectory-specific differences in memory delay. Open circles illustrate neurons without significant firing rate differences between left and right trials (DAT non-significant or Vgat non-significant). (C) Among neuronal populations, significant anatomical segregation was observed only in GABA neurons. Accordingly, across the mediolateral axis, memory-specific GABA neurons localized more lateral regions of the VTA circuit (unpaired t-test, t_(72)_ = -2.38, *P* = 0.019).

### Supplementary Table

**Table S1.**
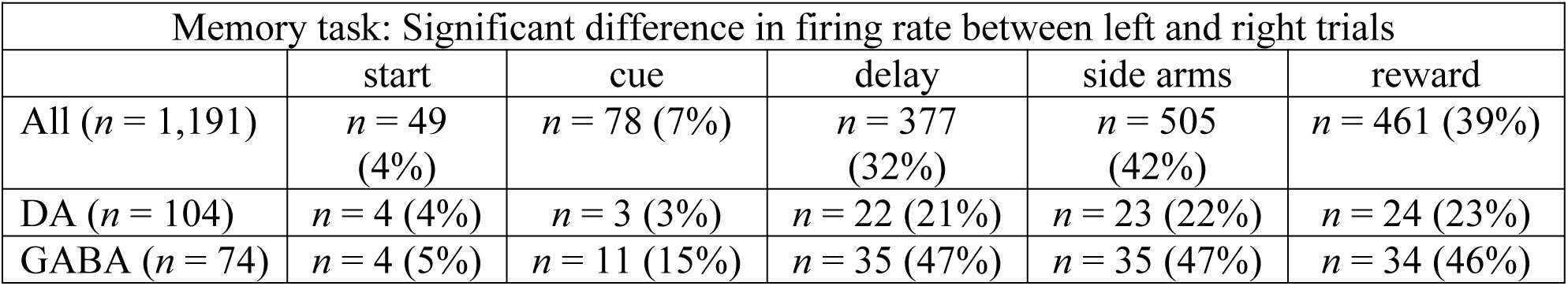
Number of neurons with trajectory-specific firing activities in the memory task grouped by maze section. Data are presented for all recorded neurons and individually for optogenetically identified DA and GABA neurons; DA, dopamine; GABA, gamma-aminobutyric acid.

## SI References

*All references have been already cited in the main manuscript*.

## References

Bäckman, C.M., Malik, N., Zhang, Y., Shan, L., Grinberg, A., Hoffer, B.J., Westphal, H., and Tomac, A.C. (2006). Characterization of a mouse strain expressing Cre recombinase from the 3′ untranslated region of the dopamine transporter locus. genesis 44, 383–390.

Berényi, A., Somogyvári, Z., Nagy, A.J., Roux, L., Long, J.D., Fujisawa, S., Stark, E., Leonardo, A., Harris, T.D., and Buzsáki, G. (2014). Large-scale, high-density (up to 512 channels) recording of local circuits in behaving animals. Journal of neurophysiology 111, 1132–1149.

Berke, J.D. (2018). What does dopamine mean? Nature neuroscience 21, 787–793.

Berridge, K.C. (2007). The debate over dopamine’s role in reward: the case for incentive salience. Psychopharmacology 191, 391–431.

Buschman, T.J., Denovellis, E.L., Diogo, C., Bullock, D., and Miller, E.K. (2012). Synchronous oscillatory neural ensembles for rules in the prefrontal cortex. Neuron 76, 838–846.

Buzsaki, G. (2006). Rhythms of the Brain (Oxford University Press).

Carr, D.B., and Sesack, S.R. (2000a). GABA-containing neurons in the rat ventral tegmental area project to the prefrontal cortex. Synapse 38, 114–123.

Carr, D.B., and Sesack, S.R. (2000b). Projections from the rat prefrontal cortex to the ventral tegmental area: target specificity in the synaptic associations with mesoaccumbens and mesocortical neurons. Journal of neuroscience 20, 3864–3873.

Choi, J.Y., Jang, H.J., Ornelas, S., Fleming, W.T., Fürth, D., Au, J., Bandi, A., Engel, E.A., and Witten, I.B. (2020). A comparison of dopaminergic and cholinergic populations reveals unique contributions of VTA dopamine neurons to short-term memory. Cell reports 33, 108492.

Cohen, J.D., Braver, T.S., and Brown, J.W. (2002). Computational perspectives on dopamine function in prefrontal cortex. Current opinion in neurobiology 12, 223–229.

Cohen, J.Y., Haesler, S., Vong, L., Lowell, B.B., and Uchida, N. (2012). Neuron-type-specific signals for reward and punishment in the ventral tegmental area. nature 482, 85–88.

Curtis, C.E., and D’Esposito, M. (2003). Persistent activity in the prefrontal cortex during working memory. Trends in cognitive sciences 7, 415–423.

Dreher, J.-C., and Burnod, Y. (2002). An integrative theory of the phasic and tonic modes of dopamine modulation in the prefrontal cortex. Neural networks 15, 583–602.

Durstewitz, D., Seamans, J.K., and Sejnowski, T.J. (2000). Neurocomputational models of working memory. Nature neuroscience 3, 1184–1191.

Engelhard, B., Finkelstein, J., Cox, J., Fleming, W., Jang, H.J., Ornelas, S., Koay, S.A., Thiberge, S.Y., Daw, N.D., and Tank, D.W. (2019). Specialized coding of sensory, motor and cognitive variables in VTA dopamine neurons. Nature 570, 509–513.

Floresco, S.B., and Magyar, O. (2006). Mesocortical dopamine modulation of executive functions: beyond working memory. Psychopharmacology 188, 567–585.

Floresco, S.B., and Phillips, A.G. (2001). Delay-dependent modulation of memory retrieval by infusion of a dopamine D₁ agonist into the rat medial prefrontal cortex. Behavioral neuroscience 115, 934.

Fries, P. (2005). A mechanism for cognitive dynamics: neuronal communication through neuronal coherence. Trends in cognitive sciences 9, 474–480.

Fujisawa, S., Amarasingham, A., Harrison, M.T., and Buzsáki, G. (2008). Behavior-dependent short-term assembly dynamics in the medial prefrontal cortex. Nature neuroscience 11, 823.

Fujisawa, S., and Buzsáki, G. (2011). A 4 Hz oscillation adaptively synchronizes prefrontal, VTA, and hippocampal activities. Neuron 72, 153–165.

Glykos, V., Whittington, M.A., and LeBeau, F.E. (2015). Subregional differences in the generation of fast network oscillations in the rat medial prefrontal cortex (mPFC) in vitro. The Journal of physiology 593, 3597–3615.

Goldman-Rakic, P.S. (1995). Cellular basis of working memory. Neuron 14, 477–485.

Goldman-Rakic, P.S. (1997). The cortical dopamine system: role in memory and cognition. Advances in pharmacology 42, 707–711.

Goldman-Rakic, P.S., Leranth, C., Williams, S.M., Mons, N., and Geffard, M. (1989). Dopamine synaptic complex with pyramidal neurons in primate cerebral cortex. Proceedings of the National Academy of Sciences 86, 9015–9019.

Hamid, A.A., Pettibone, J.R., Mabrouk, O.S., Hetrick, V.L., Schmidt, R., Vander Weele, C.M., Kennedy, R.T., Aragona, B.J., and Berke, J.D. (2016). Mesolimbic dopamine signals the value of work. Nature neuroscience 19, 117–126.

Harvey, C.D., Coen, P., and Tank, D.W. (2012). Choice-specific sequences in parietal cortex during a virtual-navigation decision task. Nature 484, 62–68.

Hauber, W. (2010). Dopamine release in the prefrontal cortex and striatum: temporal and behavioural aspects. Pharmacopsychiatry 43, S32–S41.

Howe, M.W., Tierney, P.L., Sandberg, S.G., Phillips, P.E., and Graybiel, A.M. (2013). Prolonged dopamine signalling in striatum signals proximity and value of distant rewards. nature 500, 575–579.

Jhou, T.C., Geisler, S., Marinelli, M., Degarmo, B.A., and Zahm, D.S. (2009). The mesopontine rostromedial tegmental nucleus: a structure targeted by the lateral habenula that projects to the ventral tegmental area of Tsai and substantia nigra compacta. Journal of Comparative Neurology 513, 566–596.

Kim, H.F., Ghazizadeh, A., and Hikosaka, O. (2015). Dopamine neurons encoding long-term memory of object value for habitual behavior. Cell 163, 1165–1175.

Kim, H.R., Malik, A.N., Mikhael, J.G., Bech, P., Tsutsui-Kimura, I., Sun, F., Zhang, Y., Li, Y., Watabe-Uchida, M., and Gershman, S.J. (2020). A unified framework for dopamine signals across timescales. Cell 183, 1600–1616. e1625.

Lammel, S., Hetzel, A., Häckel, O., Jones, I., Liss, B., and Roeper, J. (2008). Unique properties of mesoprefrontal neurons within a dual mesocorticolimbic dopamine system. Neuron 57, 760–773.

Lammel, S., Ion, D.I., Roeper, J., and Malenka, R.C. (2011). Projection-specific modulation of dopamine neuron synapses by aversive and rewarding stimuli. Neuron 70, 855–862.

Lee, R.S., Engelhard, B., Witten, I.B., and Daw, N.D. 2022). A vector reward prediction error model explains dopaminergic heterogeneity. bioRxiv, 2022.2002. 2028.482379.

Lee, R.S., Mattar, M.G., Parker, N.F., Witten, I.B., and Daw, N.D. (2019). Reward prediction error does not explain movement selectivity in DMS-projecting dopamine neurons. Elife 8, e42992.

Ljungberg, T., Apicella, P., and Schultz, W. (1991). Responses of monkey midbrain dopamine neurons during delayed alternation performance. Brain research 567, 337–341.

Mann, E.O., and Paulsen, O. (2007). Role of GABAergic inhibition in hippocampal network oscillations. Trends in neurosciences 30, 343–349.

Matsumoto, M., and Hikosaka, O. (2007). Lateral habenula as a source of negative reward signals in dopamine neurons. Nature 447, 1111–1115.

Matsumoto, M., and Takada, M. (2013). Distinct representations of cognitive and motivational signals in midbrain dopamine neurons. Neuron 79, 1011–1024.

Miller, E.K., and Cohen, J.D. (2001). An integrative theory of prefrontal cortex function. Annual review of neuroscience 24, 167–202.

Miller, E.K., Lundqvist, M., and Bastos, A.M. (2018). Working Memory 2.0. Neuron 100, 463–475.

Mohebi, A., Pettibone, J.R., Hamid, A.A., Wong, J.-M.T., Vinson, L.T., Patriarchi, T., Tian, L., Kennedy, R.T., and Berke, J.D. (2019). Dissociable dopamine dynamics for learning and motivation. Nature 570, 65–70.

Montague, P.R., Hyman, S.E., and Cohen, J.D. (2004). Computational roles for dopamine in behavioural control. Nature 431, 760–767.

Morris, G., Nevet, A., Arkadir, D., Vaadia, E., and Bergman, H. (2006). Midbrain dopamine neurons encode decisions for future action. Nature neuroscience 9, 1057–1063.

Moss, M.M., Zatka-Haas, P., Harris, K.D., Carandini, M., and Lak, A. (2020). Dopamine axons to dorsal striatum encode contralateral stimuli and actions. bioRxiv, 2020.2007. 2016.207316.

Nair-Roberts, R.G., Chatelain-Badie, S., Benson, E., White-Cooper, H., Bolam, J., and Ungless, M. (2008). Stereological estimates of dopaminergic, GABAergic and glutamatergic neurons in the ventral tegmental area, substantia nigra and retrorubral field in the rat. Neuroscience 152, 1024–1031.

Niv, Y. (2013). Dopamine ramps up. Nature 500, 533–535.

Omelchenko, N., and Sesack, S.R. (2009). Ultrastructural analysis of local collaterals of rat ventral tegmental area neurons: GABA phenotype and synapses onto dopamine and GABA cells. Synapse 63, 895–906.

Ott, T., and Nieder, A. (2019). Dopamine and cognitive control in prefrontal cortex. Trends in cognitive sciences 23, 213–234.

Parker, N.F., Cameron, C.M., Taliaferro, J.P., Lee, J., Choi, J.Y., Davidson, T.J., Daw, N.D., and Witten, I.B. (2016). Reward and choice encoding in terminals of midbrain dopamine neurons depends on striatal target. Nature neuroscience 19, 845–854.

Phillips, A.G., Ahn, S., and Floresco, S.B. (2004). Magnitude of dopamine release in medial prefrontal cortex predicts accuracy of memory on a delayed response task. Journal of Neuroscience 24, 547–553.

Pierce, R.C., and Kumaresan, V. (2006). The mesolimbic dopamine system: the final common pathway for the reinforcing effect of drugs of abuse? Neuroscience & biobehavioral reviews 30, 215–238.

Robbins, T.W., and Arnsten, A.F. (2009). The neuropsychopharmacology of fronto-executive function: monoaminergic modulation. Annual review of neuroscience 32, 267–287.

Salamone, J.D., and Correa, M. (2012). The mysterious motivational functions of mesolimbic dopamine. Neuron 76, 470–485.

Sawaguchi, T., and Goldman-Rakic, P.S. (1991). D1 dopamine receptors in prefrontal cortex: involvement in working memory. Science 251, 947–950.

Schultz, W. (2002). Getting formal with dopamine and reward. Neuron 36, 241–263.

Schultz, W., Apicella, P., and Ljungberg, T. (1993). Responses of monkey dopamine neurons to reward and conditioned stimuli during successive steps of learning a delayed response task. Journal of neuroscience 13, 900–913.

Schultz, W., Dayan, P., and Montague, P.R. (1997). A neural substrate of prediction and reward. Science 275, 1593–1599.

Smiley, J.F., and Goldman-Rakic, P.S. (1993). Heterogeneous targets of dopamine synapses in monkey prefrontal cortex demonstrated by serial section electron microscopy: a laminar analysis using the silver-enhanced diaminobenzidine sulfide (SEDS) immunolabeling technique. Cerebral cortex 3, 223–238.

Smiley, J.F., Williams, S.M., Szigeti, K., and Goldman-Rakic, P.S. (1992). Light and electron microscopic characterization of dopamine-immunoreactive axons in human cerebral cortex. Journal of Comparative Neurology 321, 325–335.

Stark, E., Koos, T., and Buzsáki, G. (2012). Diode probes for spatiotemporal optical control of multiple neurons in freely moving animals. Journal of neurophysiology 108, 349–363.

Stokes, M.G. (2015). ‘Activity-silent’working memory in prefrontal cortex: a dynamic coding framework. Trends in cognitive sciences 19, 394–405.

Tan, K.R., Yvon, C., Turiault, M., Mirzabekov, J.J., Doehner, J., Labouèbe, G., Deisseroth, K., Tye, K.M., and Lüscher, C. (2012). GABA neurons of the VTA drive conditioned place aversion. Neuron 73, 1173–1183.

Tobler, P.N., Fiorillo, C.D., and Schultz, W. (2005). Adaptive coding of reward value by dopamine neurons. Science 307, 1642–1645.

Traub, R.D., Bibbig, A., Fisahn, A., LeBeau, F.E., Whittington, M.A., and Buhl, E.H. (2000). A model of gamma-frequency network oscillations induced in the rat CA3 region by carbachol in vitro. European Journal of Neuroscience 12, 4093–4106.

Traub, R.D., Bibbig, A., LeBeau, F.E., Buhl, E.H., and Whittington, M.A. (2004). Cellular mechanisms of neuronal population oscillations in the hippocampus in vitro. Annu Rev Neurosci 27, 247–278.

Tsai, H.-C., Zhang, F., Adamantidis, A., Stuber, G.D., Bonci, A., De Lecea, L., and Deisseroth, K. (2009). Phasic firing in dopaminergic neurons is sufficient for behavioral conditioning. Science 324, 1080–1084.

Tzschentke, T.M. (2001). Pharmacology and behavioral pharmacology of the mesocortical dopamine system. Progress in neurobiology 63, 241–320.

Uhlhaas, P.J., and Singer, W. (2006). Neural synchrony in brain disorders: relevance for cognitive dysfunctions and pathophysiology. neuron 52, 155–168.

van Aerde, K.I., Heistek, T.S., and Mansvelder, H.D. (2008). Prelimbic and infralimbic prefrontal cortex interact during fast network oscillations. PLoS One 3, e2725.

van Zessen, R., Phillips, J.L., Budygin, E.A., and Stuber, G.D. (2012). Activation of VTA GABA neurons disrupts reward consumption. Neuron 73, 1184–1194.

Vijayraghavan, S., Wang, M., Birnbaum, S.G., Williams, G.V., and Arnsten, A.F. (2007). Inverted-U dopamine D1 receptor actions on prefrontal neurons engaged in working memory. Nature neuroscience 10, 376–384.

Vong, L., Ye, C., Yang, Z., Choi, B., Chua Jr, S., and Lowell, B.B. (2011). Leptin action on GABAergic neurons prevents obesity and reduces inhibitory tone to POMC neurons. Neuron 71, 142–154.

Wise, R.A. (2004). Dopamine, learning and motivation. Nature reviews neuroscience 5, 483–494.

